# Protocol Optimization Improves the Performance of Multiplexed RNA Imaging

**DOI:** 10.1101/2025.03.21.644527

**Authors:** Josh J. Luce, Christian A. Reardon-Lochbaum, Paolo Cadinu, Rosalind J. Xu, Jose Aceves-Salvador, Evangelia Semizoglou, Bertrand Wong, Iris Lopez, Alan B. Cantor, William Renthal, Jeffrey R. Moffitt

## Abstract

Spatial transcriptomics has emerged as a powerful tool to define the cellular structure of diverse tissues. One such method is multiplexed error robust fluorescence in situ hybridization (MERFISH). MERFISH identifies RNAs with error tolerant optical barcodes generated through sequential rounds of single-molecule fluorescence *in situ* hybridization (smFISH). MERFISH performance depends on a variety of protocol choices, yet their effect on performance has yet to be systematically examined. Here we explore a variety of properties to identify optimal choices for probe design, hybridization, buffer storage, and buffer composition. In each case, we introduce protocol modifications that can improve performance, and we show that, collectively, these modified protocols can improve MERFISH quality in both cell culture and tissue samples. As RNA FISH-based methods are used in many different contexts, we anticipate that the optimization experiments we present here may provide empirical design guidance for a broad range of methods.

## Introduction

Image-based approaches to single-cell transcriptomics have emerged as a powerful toolset for the identification and mapping of cell types and cell states in their native tissue context^1,2^. In these approaches, individual molecules are imaged by generating fluorescent signals from each targeted molecule within fixed cells and different molecules are distinguished by unique optical barcodes. In general, these optical barcodes are defined by a series of fluorescent on-off signals generated across multiple rounds of fluorescent signal generation, imaging, and fluorescence removal^1,2^. By leveraging combinatorial optical barcodes, it is possible to massively multiplex RNA imaging and simultaneously identify hundreds to tens of thousands of RNAs^1,2^. In this sense, image-based approaches to single-cell transcriptomics can be thought of as genomic-scale microscopy.

While a diversity of such methods have been introduced^1–19^, two basic classes of methods for the generation of robust, specific, and bright fluorescent signals from individual molecules have been used. In the first class, the optical barcodes for individual RNA molecules are generated by padlock probes—short DNA oligonucleotides that hybridize to the target RNA such that the 3’ and 5’ ends are adjacent and can be ligated. Once ligated these probes are amplified by rolling circle amplification, in which the circularized padlocks are amplified via a strand displacing polymerase^20^. Individual padlocks assigned to specific RNAs carry a unique nucleic acid barcode which determines the optical barcode associated with each RNA, and this barcode is read out through repetitive rounds of hybridization of complementary fluorescently labeled oligonucleotide probes^4–6,9,19^ or via sequencing chemistries^3,7,8,10^. Notable methods that use this approach to generate fluorescent signals include ISS^3^, HybISS^4,5^, SCRINSHOT^6^, STARmap^7,8^, DART-FISH^9^, BOLORAMIS^10^, and the commercial Xenium platform^19^.

In the second class of signal generation methods, modified versions of single-molecule fluorescence *in situ* hybridization (smFISH) methods are used to generate optical barcodes. In smFISH, tens of fluorescently labeled DNA oligonucleotide probes are hybridized to the sample of interest, and base pairing of these probes concentrates a large number of fluorophores at each molecular copy of the RNA, producing bright, diffraction-limited spots that can be counted to determine RNA expression levels^21,22^. As generation of large numbers of fluorescently labeled probes is expensive and hybridization of probes to cellular RNA directly is too slow for practical rounds of repetitive smFISH^11,23^, image-based approaches to single-cell transcriptomics that leverage smFISH to generate optical barcodes have instead adopted a two-step labeling process first introduced with one such method—multiplexed error robust fluorescence in situ hybridization (MERFISH)^11^. In this approach, a set of unlabeled DNA ‘encoding’ probes are created that bind to cellular RNA. These probes contain a region complementary to the RNA of interest: the ‘targeting region’; and a barcode region comprised of a series of custom binding sites: the ‘readout sequences’. The specific set of readout sequences associated with the encoding probes to each RNA determines the optical barcode associated with that RNA and is measured in successive rounds of smFISH using fluorescently labeled ‘readout probes’ complementary to each of these sequences. Hybridization of encoding probes has historically proven to be slow—typically several hours to days—while hybridization of fluorescently labeled readout probes complementary to these readout sequences is much faster^11,12^—typically minutes. Thus, this two-step labeling strategy allows rapid optical barcode readout. An important advantage of this class of methods over the previous class is that binding redundancy provided by many tens of probes to individual RNAs means that most if not all of the targeted mRNAs generate detectable signals^11^. As such, smFISH has emerged as the gold-standard for RNA copy number in single cells, and image-based approaches that use many tens of encoding probes for each RNA can have very high molecular detection efficiencies^11–14,16,17^. Notable examples of FISH-based methods are MERFISH^11–14^, seqFISH^15–17^, and the commercial CosMx^18^ and MERSCOPE platforms.

Of these methods, MERFISH may be the most adopted to date. Leveraging its combined ability to characterize large tissue areas, achieve high detection efficiencies, and profile hundreds or thousands of genes, MERFISH has driven discoveries in a wide range of tissues. Historically, neuroscience emerged as the first application space, with MERFISH being used to define and discover cell types in individual brain regions in mouse^13,24–29^ and human^30^ or within the entire mouse brain^31,32^. However, recent work has extended MERFISH to a much broader range of tissues—e.g., liver^33,34^, gut^35,36^, heart^37^, and retina^38^—and organisms, e.g., bacteria^39^ and plants^40,41^. Notably, in many cases, the discovery potential provided by the high sensitivity of MERFISH has revealed cell types and states not seen in previous studies. For example, MERFISH recently discovered a striking and unanticipated diversity of activated fibroblast states during gut inflammation^36^. In short, MERFISH, like other image-based approaches to single-cell transcriptomics, has a proven record in biological discovery.

However, many of these biological discoveries are made possible only through the high performance of MERFISH, and there are reasons to believe that further increase in performance, robustness, and experimental ease are possible. For example, properties such as the efficiency of RNA detection and the false positive rate can be set by the signal-to-noise ratio for smFISH signals. Signal brightness is set by the efficiency with which encoding and readout probes assemble onto RNAs and on the photophysical properties of fluorophores, i.e., longevity and brightness, which can be set by buffer conditions. Moreover, MERFISH measurements can extend across days and signal brightness can be modulated by the stability of reagents during this time. By contrast, background can be introduced through the generation of off-target fluorescence signals, e.g., the off-target binding of probes. Unfortunately, many aspects of encoding probe design, hybridization, buffer composition, and buffer stability have not been systematically explored; thus, it is possible that the current protocols in use^42,43^ are not ideal and that with reasonable changes improved signals can be generated.

Here we explore a variety of aspects associated with MERFISH experimental design to identify parameters that can be changed to improve performance. Specifically, we show that changes in the way in which encoding probes are hybridized to the sample can substantially improve signal brightness while modification to encoding probe design produces negligible improvements. We also show that MERFISH reagents can decrease in performance throughout the duration of an experiment, and we provide methods to ameliorate this ‘aging’ of reagents. We also systematically explore a variety of different buffers for imaging, and we introduce new buffers that can improve photostability and effective brightness for commonly used MERFISH fluorophores. Finally, we show that commonly used MERFISH readout probes can bind non-specifically in a tissue- and readout-specific fashion, that this minor increase in background can introduce false positive counts, and that this issue can be mitigated by prescreening readout probes against the sample of interest. To demonstrate the improvement provided by these protocol changes, we show improved performance for long-duration MERFISH measurements in cell culture and in colon Swiss rolls using these new protocols as compared to previously published protocols. As many aspects of the protocols optimized here are shared with other FISH-based assays, we anticipate that our improved protocols may also increase the performance of other single-RNA-molecule imaging methods.

## Results

### Signal brightness depends weakly on target region length for regions of sufficient length

FISH probes are hybridized to their target RNAs in conditions that aim to balance the conflicting goals of achieving both high assembly efficiency, i.e., the fraction of probes bound to a given RNA, and high specificity, i.e., the degree of binding to off target RNAs. This balance is typically achieved by optimizing the denaturing conditions associated with hybridization by varying a combination of temperature and the concentration of a chemical denaturant such as formamide. However, given the different inherent melting temperatures associated with target regions of different lengths, one would expect hybridization conditions to produce different results for different target region lengths. Furthermore, it is not clear that the optimal hybridization conditions associated with different target region lengths will lead to the same probe assembly efficiencies and, thus, single molecule brightness. The length of target regions varies between smFISH protocols with a typical range between 20 to 50 nt in length, yet, to our knowledge, the maximum assembly efficiency for each of these lengths has not been explored.

Thus, to explore whether there is an ideal target region length, we created a series of MERFISH encoding probe sets containing 80 different probes each with target regions that were either 20, 30, 40, or 50 nt in length. To control for potential assembly efficiency differences between different RNAs, we designed target regions to two different mRNAs (stearoyl-CoA desaturase [SCD] and chondroitin sulfate proteoglycan 4 [CSPG4]) which we anticipated would be expressed at different levels. Finally, to each of these target region sets, we affixed common readout sequences to create the encoding probes (Tables S1&S2). We then performed smFISH on U-2 OS cells with these probe sets (Methods). We identified the fluorescent signals generated by single molecules and used the brightness of these signals as a proxy for the assembly efficiency of encoding probes (Fig. 1A; Methods). As the optimal hybridization conditions will depend on target region length, we screened a range of formamide concentrations with a fixed hybridization temperature of 37 °C and hybridization duration of 1 day for each of these probe sets (Fig. 1A-C, Methods).

**Figure 1.**
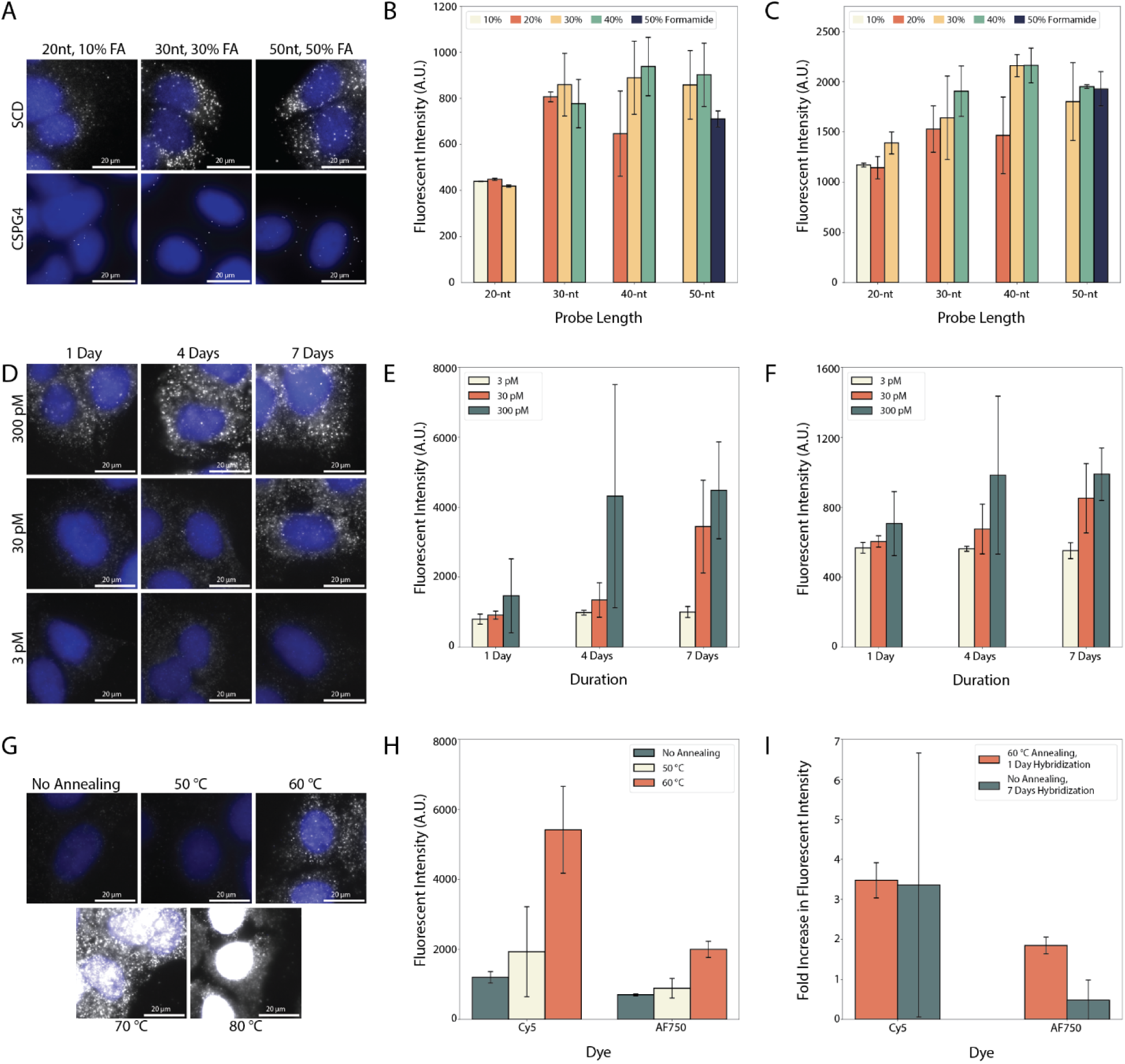
The Role of Encoding Probe Design and Hybridization on Single-Molecule Signal Brightness. (**A**) Images of U-2 OS cells stained with encoding probes that contain 20-nt (left), 30-nt (middle), or 50-nt (right) target regions and which were stained with optimal formamide (FA) concentrations for those lengths (10%, 30%, and 50%, respectively). Two different mRNAs were targeted: SCD (top) or CSPG4 (bottom). (**B**, **C**) Single-molecule signal brightness for encoding probes with different target region lengths and stained with different formamide concentrations (bar colors) for probes targeting SCD (B) or CSPG4 (C). (**D**) Images of U-2 OS cells stained with encoding probes targeting 130 RNAs and the first Cy5-labeled readout sequence. Encoding probes were stained at different per-probe concentrations (rows) and for different durations (columns). (**E**, **F**) Single-molecule signal brightness for the first Cy5-labeled readout probe (E) and the first AF750-labeled readout probe (F) for samples prepared as in (D) for the listed per-probe concentrations and hybridization durations. (**G**) Images of U-2 OS cells stained for encoding probes targeting 130 RNAs and the first Cy5-labeled readout sequence when annealed at 50, 60, 70, or 80 °C. A control without this melting and annealing protocol (No Annealing) is also shown. (**H**) Single-molecule signal brightness for the first Cy5-labeled readout probe (left) or the first AF750-labeled readout probe (right) for samples prepared as in (G) for the listed melting temperature. (**I**) The fold increase in the average molecular brightness for samples with a 60 °C melting and annealing step and 1 day of hybridization or for samples without a melting and annealing step and 7 days of hybridization as compared to a sample without melting and annealing and 1 day of hybridization. Samples were stained with an encoding probe library targeting 130 RNAs and the first readout probes conjugated to Cy5 or AF750. For A, D, and G: Gray: mRNA signal. Blue: DAPI. Scale bars: 20 µm. For B, C, E, F, H, I: Bars and error bars represent the average or standard deviation across at least two replicates of the average plotted quantities seen within each replicate.

These measurements revealed that for all probe sets the average brightness of single molecule signals depends relatively weakly on formamide concentration within the optimal range we used for each target region length (Fig. 1B&C). Moreover, these measurements suggested that signal brightness at the optimal formamide concentration depends only weakly on target region length (Fig. 1B&C). Noticeable brightness increases were seen for both SCD and CSPG4 as the target region was increased from 20 nt to 30 nt; however, within the variability of our measurements, appreciable increases were not observed as the target region length was further increased (Fig. 1B&C). These measurements indicate that once target regions are of a sufficient 30 nt length, there are limited benefits to further increases in length—at least with respect to signal brightness. Importantly, as longer target regions lead to longer encoding probes that are more costly to synthesize with existing protocols, these measurements suggest that the 30-nt target regions in current use with MERFISH^42,43^ represent an ideal choice.

### Increased hybridization duration can substantially increase probe assembly for complex encoding probe libraries

Having established the role of target region length, we next asked to what degree properties of the hybridization process can modulate single molecule brightness. Standard smFISH protocols typically stain with probe concentrations of approximately 10 nM for each probe and perform these hybridizations for 12 – 24 hours^22^. However, as the number of targeted RNAs increases in highly multiplexed measurements, maintaining a per-probe concentration of ∼10 nM becomes prohibitive. For example, targeting 1,000 RNAs with 50 probes per RNA would require 50,000 unique encoding probes and, with a 10 nM per-probe target, a total probe staining concentration of 0.5 mM. Such high probe concentrations can lead to high background, even with sample clearing methods^44^, and can be expensive. As a result, most MERFISH measurements are performed with per-probe concentrations that are at least an order of magnitude smaller, and these decreased probe concentrations are often offset with an increase in the hybridization duration to 36 to 72 hours^42,43^. However, to our knowledge it has not been established to what degree this ∼10 nM per-probe target represents a saturating concentration; thus, it is unclear to what degree decreases in per-probe concentrations or increases in hybridization durations modulate the brightness of single-molecule signals.

To address this question, we stained fixed U-2 OS cells with a previously published MERFISH encoding probe library that targets 130 RNAs with ∼90 encoding probes per RNA^12^. We stained these samples at three different total concentrations, 4 µM, 400 nM, and 40 nM, corresponding to per-probe concentrations of ∼300 pM, ∼30 pM, or ∼3 pM, and we hybridized these samples at 37 °C for 1, 4 or 7 days (Methods). To identify spots, we stained these samples with two readout probes, each conjugated to a different fluorophore, Cyanine-5 (Cy5) or AlexaFluor750 (AF750). Indeed, we found that decreasing the concentration of encoding probes resulted in a noticeable and abrupt drop in the brightness of single molecule signals for any duration of hybridization (Fig. 1D-F), indicating that these lower per-probe concentrations are, indeed, sub saturating. Importantly, increasing the duration of hybridization from 1 day to 4 or 7 days increased the brightness of spots in both color channels for some concentrations (Fig. 1D-F). Interestingly, the brightness of single molecule signals plateaued at day 4 for the highest concentration (300 pM) whereas there was a noticeable increase in brightness from day 4 to 7 for the intermediate concentration (30 pM), suggesting that 300 pM per probe is sufficient to saturate probe binding with sufficient hybridization durations. Importantly, 7 days of hybridization with 30 pM was insufficient to restore the brightness of signals observed with 300 pM, revealing either that binding is so slow that it is incomplete at day 7 or that assembly of probes is sufficient to deplete the probe in solution. In either case, this measurement suggests that 30 pM should be considered a practical lower limit for per-probe concentrations. Supporting this assertion, samples stained with 3 pM showed dim signals that failed to increase noticeably even with 7 days of hybridization (Fig. 1D-F).

### Melting and annealing can enhance the rate of encoding probe hybridization

While the previous measurements indicate that substantial increases in the duration of hybridization can improve signal brightness for modest probe concentrations, long hybridizations are cumbersome for sample preparation workflows and, critically, we noticed that the variability in brightness could be substantial with longer hybridizations (Fig. 1E&F), raising some concerns with the robustness of such approaches. Thus, we sought ways to enhance the binding of encoding probes without increasing their concentration. In this respect we were inspired by the field of DNA origami, where complex nucleic acid structures are assembled with high accuracy^45^. To aid assembly, origami protocols typically leverage an annealing process where samples are initially heated above the expected melting temperature of all oligos and then slowly reduced in temperature to favor specific binding. We reasoned that if RNA secondary structure or residual secondary structure within encoding probes were present, it may slow encoding probe binding, in which case a similar annealing protocol may accelerate binding.

To explore this possibility, we stained fixed U-2 OS cells with the MERFISH library described above at a per-probe concentration of ∼300 pM. However, once the encoding probes were applied to the sample, instead of simply incubating the sample at a fixed 37 °C, we first placed samples at 50, 60, 70, or 80 °C for 1 hour with the goal of melting secondary structure. After this initial melting incubation, we moved the samples to a 37 °C oven for a 24-hour hybridization. Importantly, the samples were transferred on a metal block that had been preheated to the temperature used in the melting step. The large heat capacity of this metal block, thus, slowed the rate at which the temperature of the block and of the samples dropped to 37 °C, providing an effective annealing process (Methods).

Remarkably, we observed a noticeable increase in the brightness of the single molecule signals relative to a protocol without this melting and annealing process for all melting temperatures other than 50 °C (Fig. 1G&H). Interestingly, we observed that the brightness of samples appeared to further increase for a melting temperature of 70 or 80 °C; however, this increase in brightness corresponded to a substantial increase in fluorescent background both in the cytoplasm and nucleus (Fig. 1G). Indeed, the samples prepared with the 80 °C protocol showed massive amounts of background fluorescence in the nucleus, consistent, perhaps, with the melting of the genome and binding of probes to single-stranded DNA (Fig. 1G). Given this increase in background, these measurements suggest that 60 °C may represent both a practical upper limit and the ideal temperature for this initial melting step under our hybridization conditions.

We next asked whether this melting and annealing process was increasing the brightness of single molecule signals by potentially unmasking binding sites occluded by secondary structure and, thus, increasing the efficiency of probe assembly or whether this process was simply accelerating the rate of encoding probe binding. To address this question, we leveraged the ability to stain with much higher per-probe concentrations when targeting individual mRNAs. We performed smFISH in U-2 OS cells with the SCD and CSPG4 encoding probes with 30-nt target regions described above with a per-probe concentration of 12.5 nM and compared the brightness of signals with and without a 60 °C melting and annealing protocol. While we did observe modest increases in the brightness of spots for both targets with melting and annealing (Fig. S1A), the increases in brightness were far more modest as compared to that observed with the lower per-probe concentrations used when 130 RNAs were targeted (Fig. 1G&H). In parallel, we asked whether a 60 °C melting and annealing protocol provided increased signal brightness relative to longer hybridizations without such treatment when samples were stained with lower per-probe concentrations associated with the targeting of 130 RNAs. Consistent with the results when only individual genes were targeted, we found that the signal brightness was comparable, if not higher, for samples hybridized for 1 day with this melting and annealing protocol as compared to those hybridized for 7 days without melting and annealing (Fig. 1I). Collectively these results suggest that most of the benefit associated with melting and annealing arises from an acceleration of hybridization rates as opposed to the unmasking of a substantial number of potential binding sites. Importantly, this protocol now makes it practical to stain samples for shorter durations with more economical encoding probe concentrations without compromising signal brightness.

### Steric hindrance does not negatively impact primary or secondary probe binding efficiency

The assembly of encoding probes on RNA is not the only hybridization that dictates the final brightness of single molecule signals. The efficiency of readout probe binding also affects these signals. Previous work has established that the binding of readout probes to encoding probes is rapid and that the 15 minute incubations we use lead to saturated binding^12^ (Methods). However, these measurements only indicate that all probes capable of binding have bound in this time; they do not rule out the possibility that because most MERFISH encoding probes contain multiple readout sequences^11,12^, the binding of readout probes to one sequence might occlude the binding of different readout probes to adjacent readout sequences via steric hinderance.

To explore this possibility, we designed encoding probes to SCD as described above but included two different readout sequences. To address potential steric hinderance, we introduced 0, 1, or 10 adenines to serve as spacers between the junction of the target region and the first readout sequence and also between the junction of the first and second readout sequences (Table S1). We stained U-2 OS samples with these probes and with the two readout probes complementary to these two readout sequences, each conjugated to either Cy5 or AF750. We found that the addition of adenine spacers did not produce a statistically significant increase in signal brightness for either channel (Fig. S1B&C). However, we did note a slight increase in the degree of non-specific binding in the cytoplasm for probes with 10 adenine spacers (Fig. S1B), which may have contributed to the appearance of a non-statistically significant increase in brightness. Thus, we conclude that steric hinderance is not a concern for readout probe binding. As the inclusion of spacers increases the length of encoding probes and, thus, the synthesis expense, we suggest that the current practice of not including spacers between readout sequences in MERFISH encoding probes is the ideal approach.

### Proper storage of readout reagents improves long-term performance

A standard MERFISH measurement involves successive rounds of readout binding, imaging, and signal removal to build optical barcodes capable of distinguishing different RNAs. The duration of a MERFISH measurement increases if the total imaged sample area is increased or as the number of targeted RNAs and, thus barcode length, is increased. For the targeting of thousands of RNAs and centimeter-squared tissue areas, it is not uncommon for a MERFISH measurement to span multiple days. It would be experimentally prohibitive to prepare the necessary buffers for each of these staining and imaging rounds prior to each round; thus, it is common to prepare these buffers at the beginning of the measurement and to store them at room temperature in an automated fluid dispensing system for the duration of the measurement^42^. However, a consequence of this approach is that reagents used for the last round of imaging may have been stored at room temperature for days prior to use. Moreover, in some fluid dispensing designs, reagent buffers are stored uncovered, open to the environment in which the experiment is conducted. We routinely use such an open-air design in our own MERFISH measurements.

To determine if there is any apparent aging to MERFISH readout reagents, we performed a three-day long MERFISH measurement targeting 130 RNAs in U-2 OS cells and compared the brightness of single molecule signals in each of the imaging rounds for both the Cy5 and AF750 channels. Indeed, we observed a clear and marked decrease in the brightness of these signals for increasing imaging rounds (Fig. 2A&B), indicating that storage at room temperature leads to a noticeable ‘aging’ of MERFISH readout reagents.

**Figure 2.**
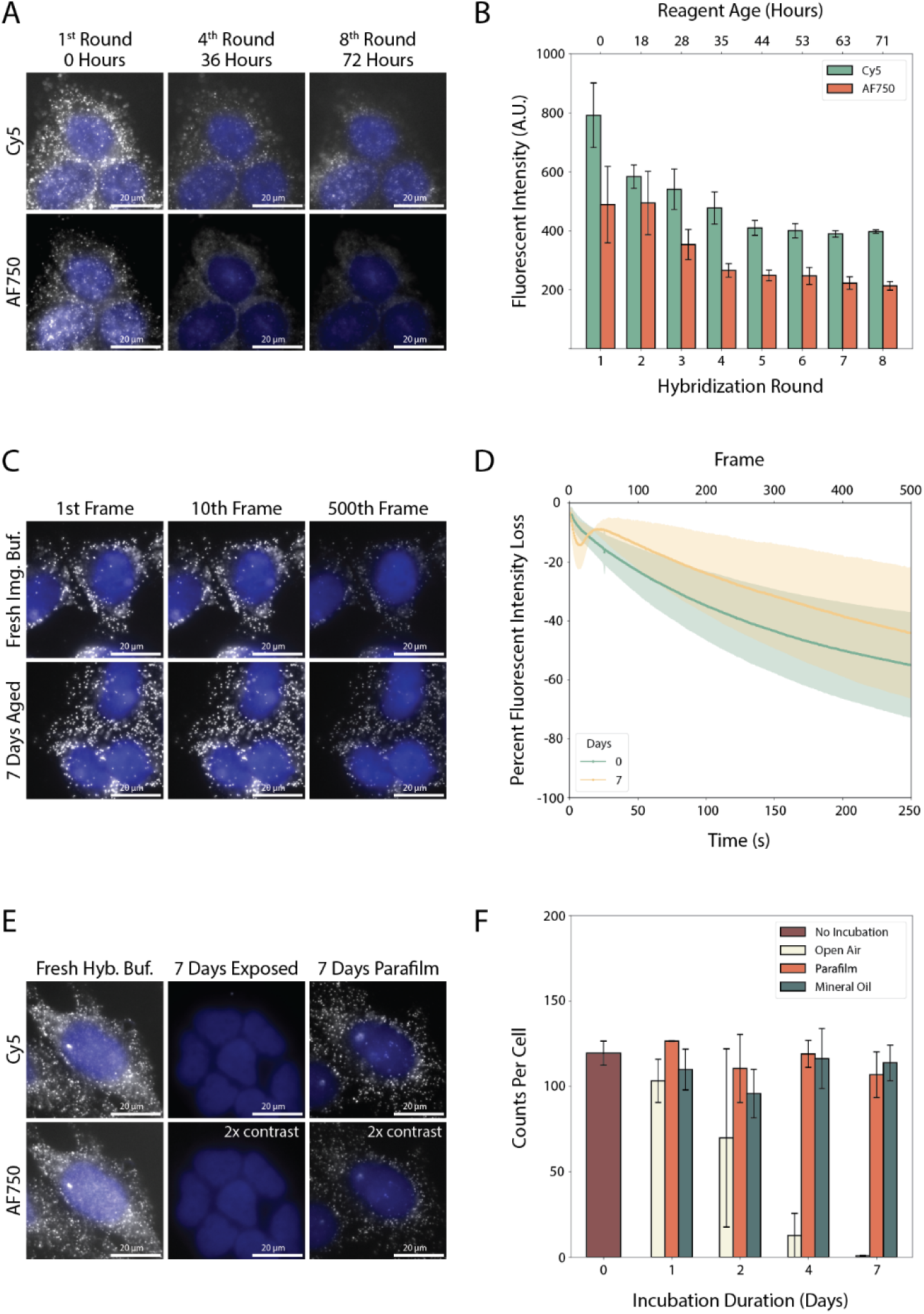
Long-Term Stability of MERFISH Readout Reagents. (**A**) Images of U-2 OS cells stained with encoding probes that target 130 RNAs and the first Cy5- or AF750-labeled readouts in the first imaging round (left), the fourth Cy5- or AF750-labeled readouts in the fourth imaging round (middle), or the eighth Cy5- or AF750-labeled readout probes in the eighth imaging round (right). The titles list the approximate time since the readout reagents were made. (**B**) Single-molecule signal brightness for the Cy5 or AF750 channels in each of the hybridization and imaging rounds with the approximate age of the readout reagents. (**C**) Images of U-2 OS cells stained with encoding probes that target FLNA and with a Cy5-labeled readout probe after 1, 10, or 500 image exposures with an imaging buffer prepared freshly (top row) or prepared and aged for 7 days at room temperature protected from oxygen (bottom). (**D**) Single-molecule signal brightness measured for the Cy5 channel versus the total exposure time for samples as in (C). Intensity is measured in precent brightness decrease relative to that observed in the first image. (**E**) Images of U-2 OS cells stained with encoding probes that target 130 RNAs and with Cy5-labeled (top) or AF750-labeled (bottom) readout probes in hybridization reagents that were prepared freshly (left) or prepared and aged in the dark but exposed to the room environment for 7 days covered (right) or uncovered (middle). Where denoted the contrast has been increased by 2-fold. (**F**) RNA copy number per cell for samples prepared as in (E). ‘No incubation’ indicates reagents prepared freshly, ‘Open Air’ indicates reagents aged in the dark but exposed to the room environment, ‘Parafilm’ and ‘Mineral Oil’ indicate reagents aged in the dark but protected from the room environment with parafilm or a layer of mineral oil, respectively. For A, C, E: Gray: mRNA signal. Blue: DAPI. Scale bars: 20 µm. For B, D, F: Bars and error bars or shaded regions represent the average and standard deviation derived from the plotted quantity from each of at least two replicates.

To obtain bright, photobleaching-resistant signals, MERFISH protocols leverage an imaging buffer^46^ that comprises both an oxygen scavenging system—which removes the dissolved oxygen that can lead to photobleaching of fluorescent dyes—and a redox pair—which reduces the lifetime of long-lived, dark triplet states by oxidizing these states and then reducing the oxidized fluorophore to its native structure. We use the common protocatechuate 3,4-dioxygenase (PCD)—protocatechuate acid system^47^ to scavenge oxygen and the Trolox-Trolox-quinone redox pair^46,48^. As multiple components of this imaging buffer could degrade with time, we first asked if degradation in the performance of the imaging buffer might explain this drop in signal brightness. To determine the rate of photobleaching of fluorophores, we repeatedly imaged the same regions of U-2 OS samples stained with encoding probe targeting the FLNA mRNA with readout sequences for a Cy5- and AF750-readout probe in either an imaging buffer prepared freshly or aged for 7 days in the dark and covered with mineral oil to prevent environmental oxygen from diffusing into the buffer (which is how this buffer is stored during MERFISH measurements; Table S1; Methods). We observed that fluorescent signals decreased as the sample was increasingly imaged, consistent with the expected photobleaching of these dyes; however, we found that both the freshly prepared and aged imaging buffer protected both Cy5 and AF750 from bleaching equally well and that the initial brightness of fluorophores was the same between the two buffers (Fig. 2C&D; Fig. S2A). This observation indicates that the imaging buffer is stable at room temperature for at least 7 days and, thus, is not the source of the drop in signal observed with time.

Having ruled out aging of the imaging buffer, we next asked if aging of the buffers associated with readout probe hybridization may drive this drop in signal. Indeed, we found that the brightness of U-2 OS samples stained with FLNA encoding probes and a Cy5 and AF750 readout probe in buffers that were aged for 7 days showed a massive reduction in signal brightness and the number of detectable RNAs per cell relative to samples stained with readout reagents prepared freshly (Fig. 2E&F, Fig. S2B-D). Thus, we conclude that aging of the readout reagents is the dominant source of the reduction in MERFISH signal brightness with time.

As readout buffers are often stored open to the environment to allow easy aspiration in our automated flow system, we next asked if contaminants in the environment might lead to the apparent aging of these buffers. To address this possibility, we aged readout hybridization buffers in the dark, as above, but covered these buffers either with parafilm or a layer of mineral oil to reduce the possibility of contamination. Remarkably, simply covering these buffers almost completely removed the apparent drop in signal brightness, even for readout reagents that were aged for 7 days (Fig. 2E&F; Fig. S2B-D). Collectively, these measurements indicate that MERFISH readout reagents are remarkably stable at room temperature, as long as they are not exposed to the environment. To address this limitation, we have modified our open-air fluid dispensing to cover readout reagents, and we recommend that when using automated fluid dispensing system that care should be taken to keep reagents covered.

### Optimized imaging buffer pH improves fluorophore photobleaching and brightness

Steady state levels of dissolved oxygen in the MERFISH imaging buffer should be low; however, when this buffer is introduced to the sample, it is possible that it is temporarily reoxygenated due to flow through components of the automated flow system. As oxygen scavenging systems require finite time to remove oxygen and residual oxygen will increase the rate at which fluorophores are bleached, we reasoned that the rate at which scavenging systems remove oxygen from buffers could produce measurable performance defects.

To explore this possibility, we first explored the degree of photobleaching observed in MERFISH samples immediately after imaging buffer was introduced to the sample via the automated flow system. We profiled photobleaching as above, and we found that there was a dramatic reduction in signal brightness in the second exposure relative to the first exposure (Fig. 3A, top) for regions of U-2 OS stained for FLNA imaged immediately after buffer was flown but not for regions imaged ∼1 minute or more after buffer flow (Fig. 3A, bottom), consistent with a transient reoxygenation of the buffer during active flow and a finite oxygen removal time by the scavenging system.

**Figure 3.**
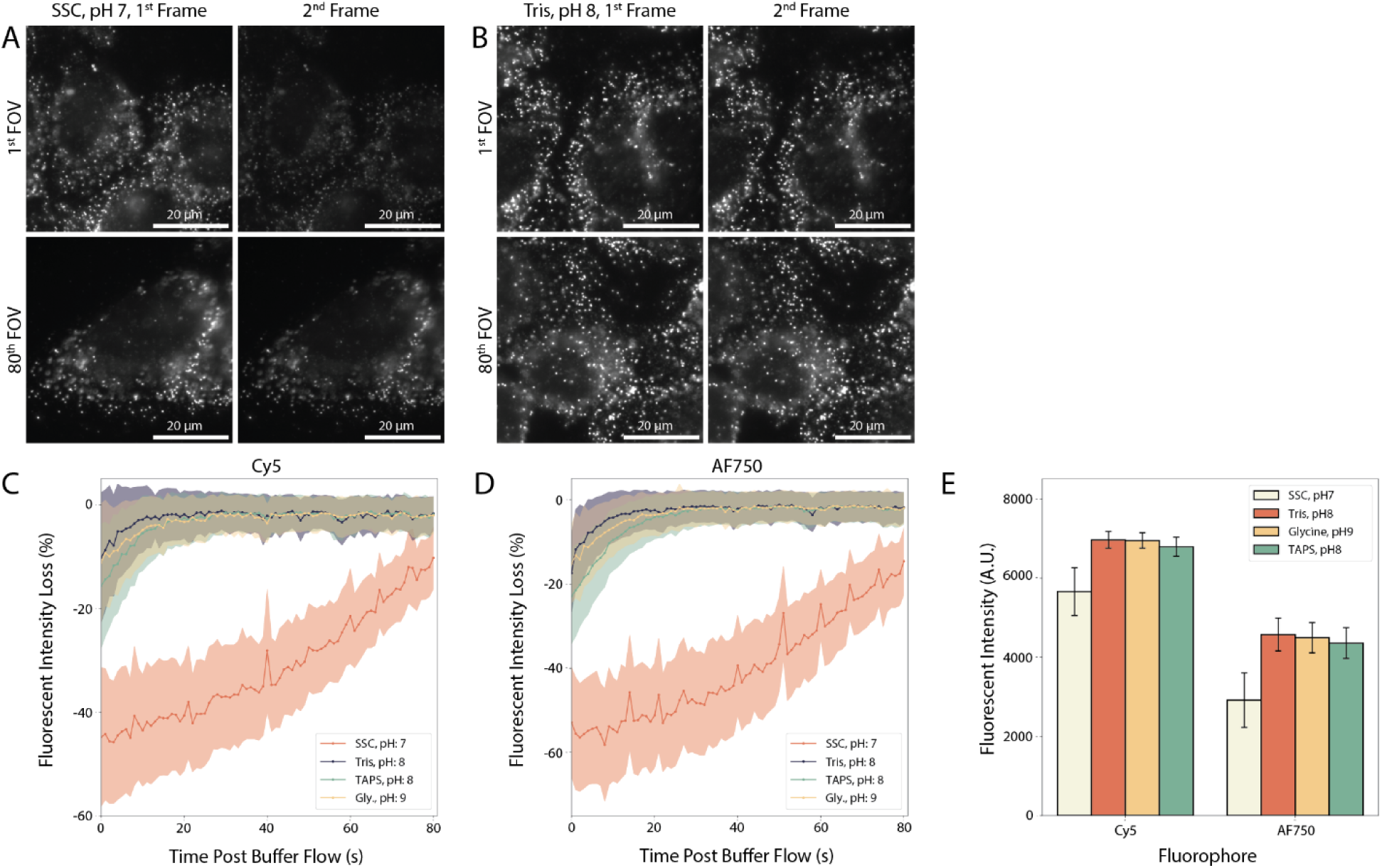
Improved Dye Brightness and Stability with Revised Readout Reagent Buffers. (**A, B**) Images of U-2 OS cells stained with encoding probes that target 130 RNAs and with the first Cy5-labeled readout probe after the first (left) or second (right) exposure of the same sample region in an imaging buffer with a base of saline sodium citrate (SSC) pH 7 (A) or a Tris-HCl pH 8 (B). The top and bottom rows represent images from the first (top) or 80^th^ (bottom) field of view (FOV) imaged immediately after imaging buffer was flown into the sample. (**C, D**) Precent loss in single-molecule brightness in the second imaged frame relative to that of the first imaged frame for Cy5 (C) or AF750 (D) imaged in imaging buffers with bases of SSC pH 7, Tris pH 8, TAPS pH 8, or Glycine (Gly.) pH 9 (Methods). (**E**) Average first-frame single-molecule signal brightness measured for the Cy5 and AF750 channels across all FOVs of the measurements in (A) and (B). For A and B: Gray: mRNA signal. Scale bars: 20 µm. For C, D, and E: Bars or solid lines represent averages and shaded areas or error bars represent standard deviation across the average of plotted quantities seen between two replicates.

While this increased photobleaching was short lived, we were concerned that it might be variable from flow system to flow system and that it could introduce artifacts into MERFISH measurements. Thus, we sought ways to remove this issue. We reasoned that increasing the activity of the oxygen scavenging enzyme, PCD, could improve the rate of recovery. The optimal pH for enzymatic activity of PCD is pH ∼9 (Refs.^49,50^), yet the imaging buffer and all other buffers associated with MERFISH have a pH of 7, which reflects the use of saline sodium citrate (SSC)—a common buffer in FISH^21,22,42,43^—as the base for these buffers. Thus, to increase the performance of the oxygen scavenging system, we explored increasing the pH of the imaging buffer. As citrate is not a good buffer at higher pH values, we replaced SSC with other more pH-appropriate buffers but kept the 300 mM sodium chloride found in SSC. Specifically, we created imaging buffers based on 50 mM Tris-HCl at pH 8, 50 mM TAPS at pH 8, and 50 mM glycine at pH 9 (Methods), and we repeated the transient oxygenation experiment above. Indeed, we found a dramatic improvement in the apparent photostability of single molecule signals immediately after imaging buffer was flown in each of these new, higher pH buffers (Fig. 3A-D), suggesting a more rapid removal of oxygen by the enhanced scavenger activity at these higher pH values. When averaged over all of the frames imaged in these experiments, switching to these higher pH buffers produced a measurable increase in the average brightness of both fluorophore channels (Fig. 3E). Moreover, at steady state, after this transient oxygenation phase had passed, we noticed an improvement in the photobleaching rate of Cy5 but not AF750 in these higher pH buffers (Fig. S3), suggesting that the enhanced activity of PCD may lower steady oxygen levels and improve the photophysical properties of some dyes. As Tris is a commonly used, inexpensive buffer, we recommend the use of Tris-based buffers at a pH of 8 for all MERFISH readout reagents (Methods).

### Sample- and readout-probe-specific background binding can increase false-positive rates

All image-based approaches to single-cell transcriptomics have the potential for the false detection of RNAs^11,14,16,17^. These false positives can be generated via a variety of mechanisms, including stray fluorescence in the sample that happens to generate patterns that match the expected optical barcodes. To estimate the rate of such false-positives, MERFISH introduced to the field the use of ‘blank’ barcodes^11^. These are valid barcodes that are not assigned to an RNA; thus, their detection provides an estimate of the rate of false positives in a measurement. Indeed, the rate of blank barcode detection provides an excellent estimate of overall false count rates in all RNAs, as we have observed that the average blank barcode level can predict the expression level below which estimates of RNA abundance determined via MERFISH are no longer correlated with those from independent techniques, such as bulk RNA-sequencing^11,13,36,39^ (Fig. 4A).

**Figure 4.**
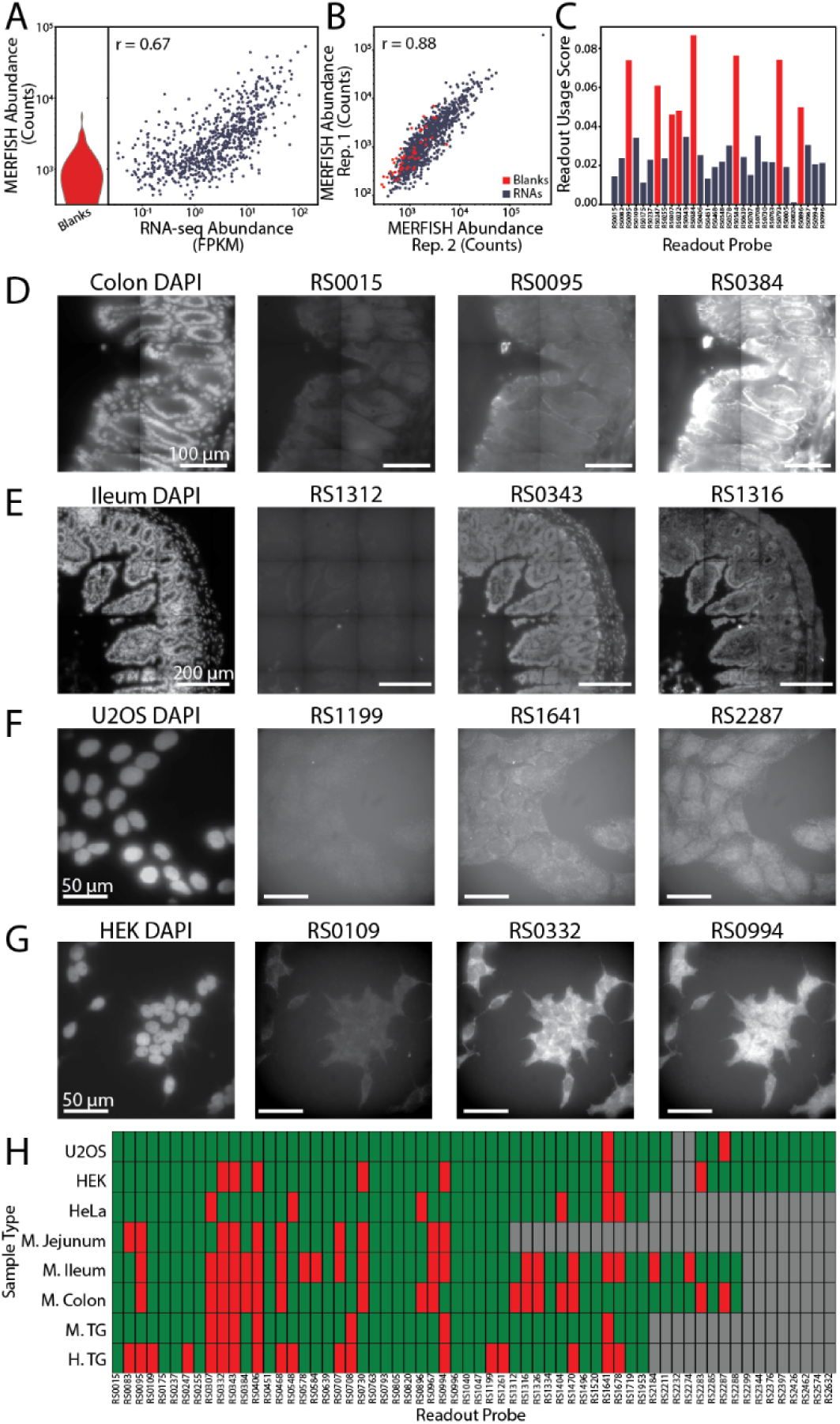
Non-Specific Readout Probe Binding Influences False Positive Rates. (**A**) mRNA abundance determined via MERFISH versus bulk RNA sequencing for measurements of the colon^36^. The distribution for the blank barcodes is included (left). (**B**) mRNA abundance determined via MERFISH from two replicates of the mouse colon^36^. mRNAs: blue. Blank barcodes: red. (**C**) Readout usage score (Methods) for all blank barcodes measured in a mouse colon dataset^36^. Red marks visible outliers. (**D-G**) Images of mouse colon (D), ileum (E), U-2 OS cells (F), and HEK293 cells (G) stained for DAPI and with the listed readout probes (Table S2). These samples were not stained with encoding probes (Methods). (**H**) Background binding score for readouts in different tissues. Red: High background. Green: minimal or no detected background. Gray: Not measured. M: mouse. H: human. TG: trigeminal ganglia. For A and B: r represents the Pearson correlation coefficient between the logarithmic expression values. FPKM: Fragments per kilobase per million reads.

MERFISH measurements typically include multiple blank barcodes, and we have noticed that the rate at which these barcodes are detected can vary between blank barcodes and, critically, that this variation is reproducible across replicate measurements^11,13,36,39^ (Fig. 4B). Thus, we asked if there were any properties that might explain why some blank barcodes are detected more frequently than others. Importantly, as such mechanisms could lead to false counts in non-blank barcodes, an understanding of their origin could reduce artifacts in MERFISH measurements. To investigate this question, we turned to published MERFISH measurements in the mouse colon^36^ and computed a ‘readout usage score’ which was defined as the fraction of blank counts that were derived from blank barcodes associated with bits assigned to each of the used readout sequences (Methods). Remarkably, we found that a subset of readout sequences had high readout usage score and, thus, were associated with high blank counts (Fig. 4C), suggesting that there may be a readout-sequence-specific origin to some of these false positive counts.

To explore why these readout sequences may be associated with increased blank barcode detection, we performed a MERFISH measurement in sections of the mouse colon that had not been stained with encoding probes (Fig. 4D). We observed that most readout probes produced very low levels of background fluorescence in these samples (e.g., RS0015 in Fig. 4D); however, there were several readout probes that produced clear patterns of staining in the absence of encoding probes, indicating that they have a propensity for non-specific binding in the mouse colon (e.g., RS0095 and RS0384 in Fig. 4D). Indeed, these high background readout probes corresponded to the readout sequences associated with a high frequency of blank barcode detection (Fig. 4C), suggesting that non-specific binding of readout probes can lead to false counts in barcodes that utilize the corresponding readout sequences. Importantly, we observed that the degree of background staining was not perfectly correlated with the detection frequency of blank barcodes that use specific readout sequences. For example, RS0384 produced much higher degrees of background staining in the mouse colon than RS0095, yet RS0384 was associated with only modestly higher degrees of blank counts as compared to RS0095 (Fig. 4C&D). Thus, we conclude that non-specific binding of different readout probes cannot be the only origin for blank barcode detection.

To test the universality of this background readout staining, we performed MERFISH on a variety of mouse tissues and several cell cultures without the application of encoding probes (Methods). Indeed, we found that across all profiled samples, we identified a variety of readout probes that bound non-specifically to the sample. Importantly, the readout probes that produced higher background were sample specific (Fig. 4D-H). For example, we observed different sets of higher background readout probes in different regions of the mouse gut (colon in Fig. 4D&H versus ileum in Fig. 4E&H). Similarly, we observed different sets of high-background readouts in different cell lines (U-2 OS in Fig. 4F&H versus HEK in Fig. 4G&H). Some readout sequences were problematic across almost all profiled tissues (e.g., RS0332 and RS0343; Fig. 4H) whereas others were problematic in only a single tissue type (e.g., RS0578; Fig. 4H). Importantly, the background binding also varied by organism even within the same tissue (e.g., RS0083 in human versus mouse trigeminal ganglia [TG]). Interestingly, none of the readouts used with the first 16-bit MERFISH library that introduced these readout sequences^12^ were high-background readout sequences in the cell line in which they were first demonstrated (U-2 OS; Fig. 4H), providing an explanation for why this effect was not originally characterized. In summary, we conclude that non-specific readout probe binding is sample specific and that it can lead to elevated rates of false positives for barcodes that use those readout sequences. For this reason, we recommend that MERFISH readout probes be screened against each sample of interest and that only readout sequences associated with low-background readout probes be used to generate barcodes for those samples.

### Optimized MERFISH protocols improve performance in long-duration cell culture measurements

We next asked whether these improved protocols could improve the performance of MERFISH measurements. To explore this possibility, we performed MERFISH measurement of 130 RNAs in U-2 OS cells. We prepared and ran two biological replicate samples each using either the published MERFISH protocols or the optimized protocols developed above. Namely, in the optimized protocols we incorporated the new imaging and readout buffers, a 60 °C melting and annealing step, and we leveraged a modified automated fluidics system design that covers readout reagents (Methods). As the 16 readout sequences used previously were associated with low-background readout probes in U-2 OS, we kept the same MERFISH encoding probes used previously^12^. To illustrate the enhanced long-term performance of our measurements, we imaged sufficient area to require 3 days per measurement.

Indeed, we found that the quality of the single-molecule signals in the first round of imaging were noticeably improved with the optimized protocols as compared to that seen with the previous protocols (Fig. 5A, left). Moreover, the difference in brightness was even more striking when images from the final round of single-molecule imaging were examined (Fig. 5A, right), consistent with the substantial aging of readout reagents with previous methods. These brightness differences were also reflected in the quality of the decoded MERFISH data. Samples run with the previous protocols had far fewer RNAs identified per cell than samples run with the optimized protocols (Fig. 5B). In addition, the quality metrics associated with identified RNAs were also substantially higher with the optimized protocols. For example, one such quality metric is the aggregate brightness of RNA molecules across all imaging rounds (Methods). When we compared the histogram of these brightness values between these measurements, we noticed that while both had a population of low brightness molecules, which represent noise in the decoding process that is routinely filtered with a brightness threshold (dashed line in Fig. 5C), the optimized protocols produced molecules that were substantially brighter than those identified with the old protocols (Fig. 5C). Thus, we find that the optimized protocols identify more RNAs with higher quality metrics than the old protocols.

**Figure 5.**
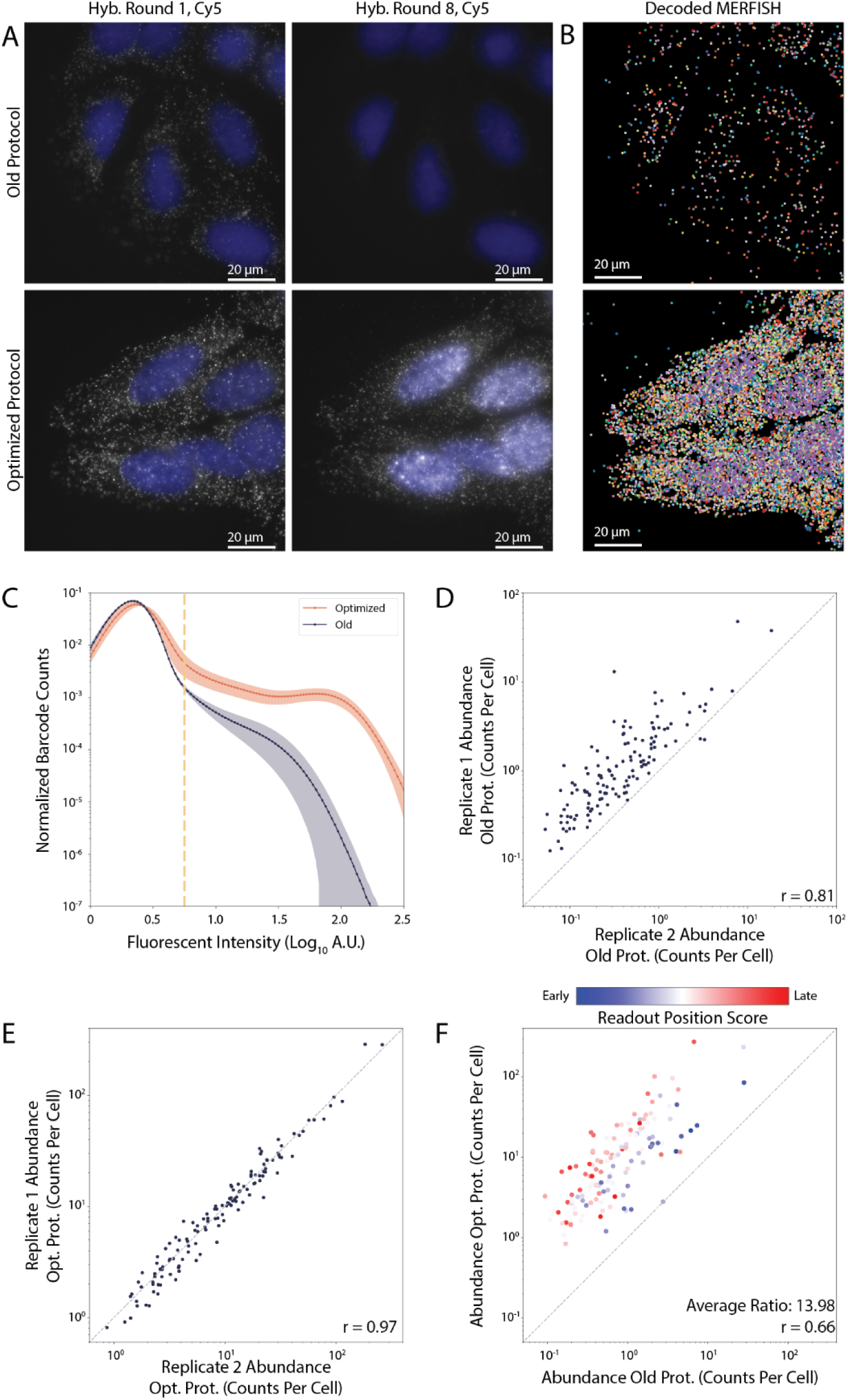
Optimized Protocols Improve MERFISH Performance in Long-Term Measurements of Cell Culture. (**A**) Images of the first (left) or last (right) hybridization round of MERFISH U-2 OS cells for the Cy5 channel run with the published protocols (top) or the optimized protocols presented here (bottom). (**B**) Location and identity (color) of all identified RNAs from the MERFISH measurements in (A). (**C**) Histogram of molecular brightness—an important quality metric for identified RNAs—for all molecules identified with the previous (blue) and optimized (red) protocols. The dashed line is the threshold used to discriminate high from low quality RNAs. (**D**, **E**) The average RNA copy number per cell for all genes versus that of another measurement for two replicate measurements with the old protocols (D) or with the optimized protocols (E). (**F**) The average copy number determined from the average of both MERFISH replicates with the optimized protocols (Opt.) versus that of the published protocols (Old). mRNAs are colored by the readout position score, with red indicating a readout determined by later hybridization rounds. For D, E, and F: the dashed line is equality. r represents the Pearson correlation coefficient between the logarithmic expression values.

This improvement in quality was also reflected in measures of reproducibility. We found there was greater variability in the average RNA copy number per cell measured between the two replicate measurements with the old protocols (Fig. 5D) than there was between the two replicates produced with the optimized protocols (Fig. 5E). To explore potential origins for this increased variability, we compared the average RNA copy number per cell between the two protocol sets (Fig. 5F). This comparison confirmed that, on average, we detect ∼14-fold more RNA copies per cell with the new protocols relative to the old protocols (Fig. 5F). Nonetheless, we observed substantial variability between the measurements of the two protocols with some RNAs detected far more frequently than others with the new protocols relative to the old (Fig. 5F). We reasoned that this variation may arise from systematic variations in the efficiency with which RNAs are detected based on the specific imaging rounds associated with their barcodes. To explore this possibility, we created a readout position score—defined as the sum of the imaging rounds in which the barcode has ‘1’ values (Methods). Barcodes that leverage readout sequences associated with later imaging rounds would have larger readout position scores while barcodes that leverage readout sequences associated with earlier imaging rounds would have smaller scores. Remarkably, we find that much of this variability between the abundance measurements with the two protocol sets can be explained via this readout position score (Fig. 5F). This observation is consistent with a systematic variation in the detection efficiency with the old protocols: RNAs with barcodes that leverage latter imaging rounds are detected less frequently, likely reflecting the dimmer signals in these rounds. It is important to note that the lower performance we observe with the old protocols is not reflective of what has been reported previously^12^, as care was taken in previous measurements to ensure that measurements did not extend for the long durations used here. These measurements were designed, instead, to illustrate the increased long-term performance of the optimized protocols.

Taken together, these measurements reveal that the three key optimizations that we introduced here—melting and annealing to improve encoding probe hybridization, Tris-based pH 8 readout reagent buffers, and careful storage of readout reagents to prevent aging—improved RNA detection efficiency, increased molecular brightness, and decreased barcode-specific artifacts in long MERFISH measurements in cell culture. For these reasons, we recommend these new protocols for all such measurements.

### Optimized MERFISH protocols improve performance in large tissue samples

We next asked whether the optimized protocols can improve performance in tissue imaging. To this end, we harvested the colon of a C57BL/6J wild-type mouse and created a fixed-frozen Swiss roll of the entire colon (Methods). We then stained this sample with a published MERFISH library targeting 940 RNAs^36^, and we designed and stained a revised version of this library that excluded the readout sequences that we identified as potentially problematic in the mouse colon (Fig. 4; Table S3&S4; Methods). For our measurement with the published library, we leveraged the old protocols. Specifically, we used SSC buffers and performed a 3-day long encoding probe hybridization without a melting and annealing step. For our measurements with this new library, we leveraged our optimized protocols, including the new Tris-based buffers at pH 8 and a 60 °C melting and annealing step followed by a 1 day encoding probe hybridization. Measurement of the entire Swiss roll in both cases required less than two days; thus, the total measurement time was not as long as that used for the cell culture measurements above.

The MERFISH measurements for both samples were of sufficient quality to resolve the structure of the colon through the distribution of markers of anatomical features (Fig. 6A&B). The epithelial layer was marked by Vil1, the stem cell niche by Mki67, the stromal compartment by Col1a2, the muscle layers by Tagln, the lymphatic vessels in the sub-mucosa by Lyve1, the enteric nervous system by Hand2, and B cells in the lamina propria and gut associated lymphoid tissues by Cd79a (Fig. 6A&B). Moreover, we observed the total abundance of mRNAs were highly correlated between the two measurements (Fig. 6C).

**Figure 6.**
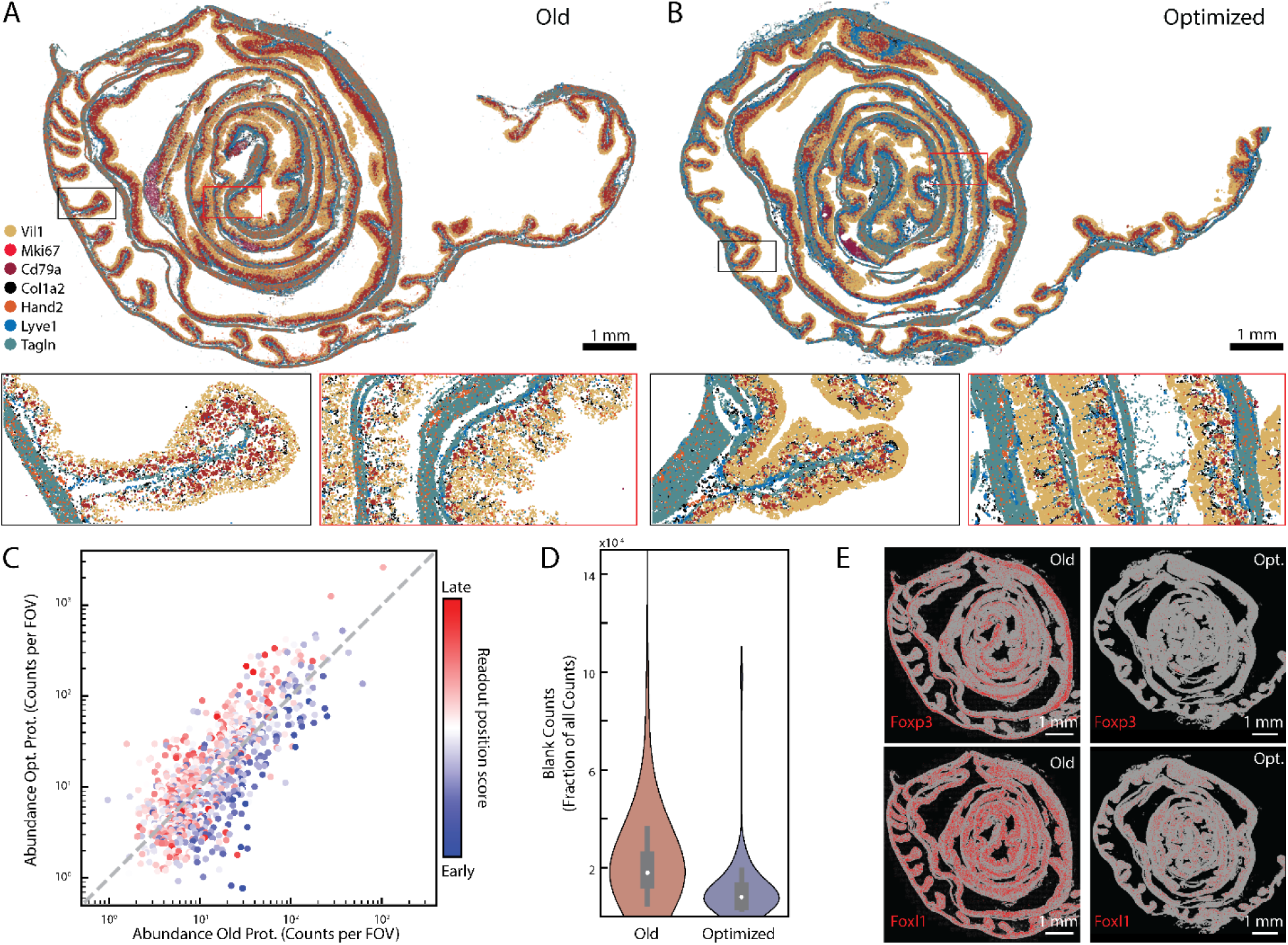
Optimized Protocols Improve MERFISH Performance in Tissue Samples. (**A, B**) The spatial distribution of 7 out of ∼1000 mRNAs profiled in a Swiss roll of the mouse colon using a published library and published protocols (A) or using a new library that removes high-background readout probes and the optimized protocols presented here (B). Zoom-ins are boxed. (**C**) mRNA abundance determined with the new library and optimized protocols as in (B) versus that measured with the published library and protocols in (A). mRNAs are colored by readout position score (Methods). (**D**) Distribution of the abundance of blank barcodes measured in fraction of all detected barcodes. (A,B). (**E**) Spatial distribution of Foxp3 (top) or Foxl1 (bottom) in the measurement in (A; left) or (B; right). All mRNAs are plotted in gray and these two mRNAs are plotted in red. For A,B: Scale bars: 1 mm.

While we did not see the same dramatic improvement in mRNA detection with the optimized protocols relative to that seen with the old protocols as observed with cell culture (Fig. 5F vs Fig. 6C)— likely reflecting the shorter duration of these measurements relative to those in cell culture—we, nonetheless, observed a reduction in artifacts associated with reduced signal quality in later imaging rounds as compared to the old protocols. To illustrate this point, we first computed the readout position score, as above, and again found that it partially explained the variation in abundance observed for some mRNAs measured with the old protocols relative to the new protocols (Fig. 6C; Methods). These measurements suggest that even over relatively short duration measurements, the reduced brightness of signals in later imaging rounds disfavors the detection of mRNAs that leverage signals in these rounds for these barcodes. However, by reducing this decrease in brightness, the optimized protocols remove this modest source of bias.

We also found that there was a reduction in the false positive rates with the optimized protocols relative to the old protocols, as judged by the frequency with which the blank barcodes were detected (Fig. 6D). This reduction in false positive rates can have important consequences for the biological interpretation of these data. To illustrate this point, we explored the distribution of two important markers of populations in the gut: Foxp3 which marks T regulatory cells and Foxl1 which marks fibroblasts in the mucosa. In previous work with the published colon MERFISH library^36^, we noted aberrantly high expression of both of these key markers, as well as expression in tissue locations and cell types for which there were no prior reports of either Foxp3 or Foxl1 expression. As these observations suggested that these genes were contaminated by false positive counts, we did not draw any biological conclusions from their measurements^36^. Importantly, Foxp3 and Foxl1 were encoded with barcodes that leveraged readout sequences consistent with high-background readout probes (RS0384 and RS0332 in Fig. 4F), consistent with a readout-background-induced elevated false positive rate. We reproduced these artifacts in the Swiss roll measurements here with the published library and protocols (Fig. 6E). Importantly, with the new protocols and the new library, in which high-background-readout probes were not used, the problematic distribution of these two important markers is no longer seen (Fig. 6E). This observation underscores the potential benefits of the lower false positive rates provided by prescreening the background of readout probes in tissue of interest and then designing MERFISH encoding probes that use the readout sequences associated with low-background readout probes.

Collectively, these measurements reveal that our optimized protocols increase multiple aspects of MERFISH performance in tissue. When taken together with the performance improvements we observed for long-duration measurements in cell culture, we, thus, recommend these new protocols for all MERFISH measurements.

## Discussion

Massively multiplexed single-molecule RNA imaging has emerged as a powerful new approach to the definition and charting of cell types and cell states in a wide variety of contexts^1,2^. Here we examined a variety of protocol choices in one such method, MERFISH^11,42,43^, and identified a variety of changes that improve the performance of this technique in cell culture and tissue. Importantly, we provide guidelines based on these protocol improvements to aid other experimentalists in the design of their own MERFISH measurements.

We first examined the properties of MERFISH encoding probe design. FISH probe length varies broadly in the literature^11,15,21,22^; yet there has not been a systematic side-by-side comparison of the assembly efficiency of FISH probes of different length. Our measurements indicate that it is possible to obtain brighter signals by increasing the length of target regions from 20 nt to 30 nt but beyond 30 nt the benefits to brightness, if they exist, are modest. As target regions of 20 nt length remain a common choice for conventional, singleplex smFISH^22^, we would propose that these measurements may benefit slightly by an increase in target region length. By contrast, the original MERFISH protocols introduced 30 nt target region lengths^11^, and this length remains the most common choice^12,13,24–28,30,31,33–36,38^. Thus, we do not believe there is a need to change these practices. We also explored the potential for steric hinderance between the binding of target regions to their target RNA and readout probes or between the binding of one readout probe with that of readout probes targeting adjacent readout sequences and find no evidence that this type of hinderance is a concern. Thus, there is no need to incorporate extra nucleotides as spacer elements into the design of MERFISH encoding probes.

Hybridization is also a critical determinant of encoding probe assembly efficiency. Multiplexed measurements typically use small per-probe concentrations of encoding probes relative to that in standard singleplex measurements^22,42,43^. We now show that this decrease in concentration does, indeed, come at a cost in the rate at which encoding probes bind to the sample, and we provide a practical lower limit to the per-probe concentration of ∼30 pM. As multiplexed FISH measurements target increasing numbers of mRNAs, this practical lower limit may guide target staining concentrations. Moreover, we show that increased hybridization durations can, at least partially, offset this lower per-probe concentration. However, we recommend instead that users consider the melting and annealing protocol we introduce here, as we find that it produces comparable single-molecule brightness to that of multi-day-long hybridizations in far shorter times and with less variability in replicate-to-replicate brightness. Moreover, as multi-day preparation protocols can impede rapid testing and experimental progress, we anticipate that the reduction from multi-day to single-day hybridizations may prove very convenient.

We also addressed an observation regarding the stability of MERFISH readout reagents. Importantly, we showed that reagents age over the span of days, producing measurable decreases in the quality of single-molecule signals and introducing artifacts into the detection of different RNAs by MERFISH. Interestingly, we found that all MERFISH readout reagents are remarkably stable, as we see no performance defects in either the imaging buffer or the readout hybridization reagents when stored at room temperature for a week, as long as they stored protected from the lab environment. We now show that our previous practice of open-air storage of these reagents, while convenient for automated flow systems, led to contamination from the laboratory environment. While we have not identified the contaminants—we suspect nucleases that led to the degradation of readout probes—we show that covering these solutions is a straightforward and simple solution to this problem.

In parallel, we explored the composition of buffers used for MERFISH readout. The original composition of these buffers was based on SSC, which has been a longstanding choice in the FISH community. However, we found that the pH of this buffer is not well matched to the performance of the oxygen scavenging system we use to increase fluorophore brightness and photostability. While there are other enzymatic oxygen scavenging systems that could be utilized with MERFISH^51^, PCD is a particularly attractive choice given its cost, modest effect on the pH of the buffer as oxygen is scavenged^51^, and its remarkable stability at room temperature for long durations^49,50^. Thus, we sought instead to align the buffers used with MERFISH to the properties of PCD. Indeed, we found that simply increasing the pH from 7 to 8 made a noticeable improvement in the rate at which PCD removed oxygen from solution and that steady state fluorophore brightness increased, albeit slightly, from a likely drop in the steady level of dissolved oxygen. Based on these measurements, we recommend switching all MERFISH readout reagents from SSC pH 7 to Tris pH 8.

Finally, we discovered that modest degrees of off-target binding by MERFISH readout probes can produce fluorescent background that leads to an increased rate of false positives for barcodes that leverage those readout probes. Interestingly, the non-specific readout binding is tissue, cell-type, and organism specific. While we have not determined the molecular targets of this non-specific binding, given that our clearing methods remove many proteins and lipids, it is highly likely that this off target binding is to other cellular RNAs. It may be possible to develop improved readout hybridization approaches that increase the specific of readout probes; however, we show here that a simple and straightforward solution is to prescreen the sample of interest and then design MERFISH encoding probes such that problematic readouts are excluded. Moreover, we show that such screening can reduce false positive rates in tissue. A lower false positive rate, in turn, is particularly important when one considers the biological interpretation of these measurements. More broadly, this observation underscores how important it is for users of this new suite of image-based spatial transcriptomics methods to recognize that technical details of how these measurements are performed can modify artifacts that must be kept in mind as they interpret this new class of data.

Importantly, we show that these collective protocol improvements can have important consequences for the quality of MERFISH measurements, and we demonstrate performance increases with these optimized protocols relative to published protocols for both cell culture and tissue measurements. Image-based approaches to single-cell transcriptomics, such as MERFISH, offer exciting new windows into the diversity and organization of cells within tissues and of the organization of the transcriptome within cells in vast array of tissues, cell lines, and organisms. Thus, we anticipate that the improved protocols and the higher quality data they engender will empower a wide variety of exciting discoveries in a wide range of biological contexts.

## Acknowledgments

We thank members of the Moffitt laboratory for helpful discussions. We thank R. Shim, J. Hurley, and S. Aviles for initial conversations and pilot measurements that led to the optimization efforts described here. Portions of this research were conducted on the O2 High Performance Compute Cluster, supported by the Research Computing Group, at Harvard Medical School. This work was funded through National Institutes of Health grants (R01GM143277 to JRM; U19NS130617 to JRM and WR; R01HL159106 to JRM and ABC; and 5T32HL007574-36 which supported CAR-L), from a grant from the Mathers Foundation (to JRM), and by the Charles A. King Postdoctoral Research Fellowship Program from the Bank of America Co-trustees (to PC).

## Contributions

JJL and JRM conceived the project. JJL performed all optimization experiments. JJL and CAR-L analyzed optimization experiments. CAR-L, PC, RJX, JAS, BW, and ES performed readout probe screening measurements. IL and ES provided human TG. PC performed Swiss roll measurements. JJL, CAR-L, PC, and JRM wrote the manuscript draft. JJL, CAR-L, PC, RJX, JAS, ES, BW, IL, WR, ABC, and JRM revised the manuscript draft. WR, ABC, and JRM provided supervision and funding.

## Competing interests

JRM is a co-founder of, stakeholder in, and advisor for Vizgen, Inc. JRM is an inventor on patents associated with MERFISH applied for on his behalf by Harvard University and Boston Children’s Hospital. JRM’s interests were reviewed and are managed by Boston Children’s Hospital in accordance with their conflict-of-interest policies. CAR-L and PC are inventors on patents associated with aspects of MERFISH not included in this work applied for on their behalf by Boston Children’s Hospital.

## Materials and Methods

### smFISH and MERFISH probe design and production

Single molecule FISH probes were designed against the genes SCD and CSPG4 with varying target region lengths of 20, 30, 40, and 50 nucleotides. These genes were selected because they have single isoforms, are long enough to support sufficient unique target regions, and vary in expression by ∼10-fold. Potential target regions were designed as described previously^12^ using the same pipeline, the GRCh38 human transcriptome, and publicly available RNA sequencing data for this cell line (GEO GSM1231610)^52^. Briefly, probes were screened for the predicted melting temperature (± 7.5 °C around the median value for all potential probes), and for GC content (± 15% around the median value for all probes), with the same constraints applied to target regions of varying length. 80 unique target regions were randomly selected for each gene, and encoding probes were designed by concatenating the readout sequence associated with RS0015 to each with a single adenine spacer as has been used previously (Tables S1&S2). Probes to explore the role of steric hinderance (Fig. S1) were designed as described above with two notable differences. First, each encoding probe contained a readout sequence for both RS0015 and RS0083. Second, a variable number of adenine bases were added between the target region and the first readout sequence and between the first and second readout sequence to serve as spacers. Target regions for FLNA smFISH were taken from previously published sequences^11,12^. These target regions were then concatenated with readout sequences RS0015 and RS0083. All smFISH encoding probes (Table S1) were directly synthesized by IDT. smFISH probes were column purified (Zymo, D4004) prior to use.

The 940-RNA MERFISH encoding probe set used to target mouse colon after readout probe screening was designed using previously described methods^12^. Briefly, we leveraged the GRCm39 build of the mouse transcriptome and estimated RNA abundances using previously published bulk RNA-sequencing data (GEO: GSE229027)^36^. We designed probes with a threshold of 0.75 – 1 for their gene specificity, 65 – 75 °C for their predicted melting temperature, and 0.4 – 0.6% for their GC content. Where possible, probes were designed to be isoform specific leveraging an isoform specificity range of 0.75 – 1. Target regions were allowed to overlap by as much as 20 nt. Readout sequences were selected by prescreening the background binding of readout probes and selecting only sequences associated with low-background probes (Fig. 4). A 30-bit Hamming Weight 4, Hamming Distance 4 barcoding scheme was used to encode these RNAs, leaving 50 barcodes to serve as ‘blank’ controls. 72 target regions were designed per gene with the exception of a small number of genes which were not of sufficient length to support this number of unique target regions. In these cases, the maximum number of target regions possible were used. Encoding probe template molecules were then constructed by concatenating 3 of the 4 readout sequences associated with each RNA to each target region and affixing a forward primer sequence and the reverse complement of the T7 promoter (both needed for synthesis of probes). The 130-RNA MERFISH encoding probe set used with U-2 OS cells was published previously^12^. The mouse colon 940-RNA MERFISH library that did not incorporate readout probe screening was also previously published^36^.

Template oligopools for these encoding probes were synthesized by Genscript (130-RNA U-2 OS MERFISH encoding probe set) or Twist Biosciences (for all others). Large quantities of single-stranded DNA encoding probes were amplified using a previously published amplification protocol that involves qPCR, *in vitro* transcription, reverse transcription, and alkaline hydrolysis^43^. All necessary primers were ordered from IDT.

### Cell culture and preparation

U-2 OS (ATCC, HTB-96) cells were cultured in McCoy’s 5A Modified Medium (Thermo, 16600082) containing 10% (v/v) fetal bovine serum (FBS; Fisher, MT35010CV) and were plated directly on 40 mm coverslips (Bioptechs, 40-1313-03193). Prior to use in cell culture, coverslips were silanized as previously described^43,44^. Cells were cultured at 37 °C in 5% CO_2_ for 24-72 hours to achieve a cell density of ∼1,600,000 cells per coverslip. Cells were fixed and permeabilized as described previously^12^. Briefly, cells were fixed to coverslips with 4% (v/v) paraformaldehyde (PFA; EMS, 15714) in 1× phosphate buffered saline (PBS) (Invitrogen, AM9625) for 20 minutes at room temperature. Coverslips were washed 3 times in 1× PBS, then permeabilized in 0.5% (v/v) Triton-X (Sigma, T8787) in 1× PBS for 10 minutes at room temperature, or were permeabilized in 70% ethanol (Fisher, 22-032-600) at 4 °C overnight. Fixed cells were stained immediately or stored in 70% ethanol at 4 °C up to 7 days. Human embryonic kidney (HEK 293; ATCC, CRL-3216) cells were cultured in Dulbecco’s Modified Eagle’s Medium (DMEM; Sigma, D5796) containing 10% (v/v) FBS and were otherwise prepared identically to U-2 OS. HeLa cells were cultured in DMEM containing 10% (v/v) FBS and 0.2% (v/v) Penicillin-Streptomycin (Gibco, 15140122) and were otherwise prepared identically to U-2 OS.

### Tissue harvest and preparation

Mouse gut tissues were harvested from 14 weeks old C57BL/6J male mice from Jackson Labs (Strain #000664) as described previously^43^. Mice were housed in the Harvard Center for Comparative Medicine (HCCM) facility. All mouse experiments were performed in compliance with NIH guidelines and were reviewed and approved by the Harvard Institutional Animal Care and Use Committee (IACUC) under protocol IS00003215 (all mouse gut regions) or by the Mass General Brigham IACUC under protocol 2020N000110 (mouse trigeminal ganglia).

For colon Swiss rolls, mice were euthanized with isoflurane (Patterson, 07-890-8115) followed by cervical dislocation. The colon was immediately dissected out, cut open and rinsed with 4% v/v PFA in 1× PBS at 4 °C to remove the feces and then fixed with 4% (v/v) PFA in 1× PBS at 4 °C for 48 hours. The tissue was then dehydrated in 30% (w/v) sucrose (Fisher, 419762500) in nuclease free water overnight at 4 °C. Colon was then rolled into a Swiss roll and then embedded in OCT (Fisher, 1437365), frozen on dry ice, and blocks were stored at -80 °C until sectioning. Prior to sectioning, silanized coverslips were treated with poly-D-lysine (Gibco, A38904-01) and fiducial beads (Thermo, F13082) to aid in tissue adherence and align MERFISH imaging rounds. Cryoblocks were sectioned with a cryostat at 10-µm thickness and tissue slices were melted onto coverslips. After sectioning, tissue slices on coverslips were dried at room temperature for 30 minutes, fixed with 4% (v/v) PFA in 1× PBS, washed twice with 1× PBS, and permeabilized overnight in 70% ethanol at 4 °C. Samples were either stained the day after or stored at 4 °C for up to 7 days.

For all gut regions profiled for readout probe screening, mice were euthanized using isoflurane followed by cervical dislocation. Distinct sections of the intestine were rapidly dissected, flushed with 4 mM ribonucleoside vanadyl complex (NEB, S1402S) in 1× PBS at 4 °C, and embedded in OCT. The tissue was frozen on dry ice and stored at -80 °C until sectioning. Tissue cryoblocks were sectioned at 10-µm thickness, melted onto coverslips coated with poly-D-lysine (Gibco, A38904-01) and fiducial beads (Thermo, F13082), dried for 2 hours at -20 °C inside the cryostat to improve adherence, and then fixed in 4% PFA in 1× PBS for 20 minutes at 4 °C. The tissue was washed twice with 1× PBS and permeabilized overnight in 70% ethanol at 4 °C. Samples were either stained the following day or stored at 4 °C for up to 7 days.

Mouse trigeminal ganglia were dissected from 8–10-week-old mice as described previously^53^. Trigeminal ganglia were cryoprotected in 30% w/v sucrose and the mounted flat in OCT to maintain the V1, V2, and V3 branch orientation. Tissue was then sectioned at a 14-µm thickness onto coverslips coated with poly-D-lysine and fiducial beads, dried for 2 hours at -20 °C inside the cryostat, and then fixed in 4% PFA in 1× PBS for 20 minutes at 4 °C. The tissue was washed twice with 1× PBS and permeabilized overnight in 70% ethanol at 4 °C.

Human trigeminal ganglia procurement was conducted under protocol MGB1924, which was reviewed by the Institutional Review Board (IRB) of Mass General Brigham and determined to not be restricted human research. Subjects included in this study were adults, whose autopsy started close to the time of death and were negative for human immunodeficiency virus (HIV), hepatitis B virus (HBV), hepatitis C virus (HCV), or suspected of Creutzfeldt Jacob’s disease (CJD). Following brain removal, the trigeminal crescent, rootlets, and trigeminal nerve along the medial aspect of the petrous ridge in the middle of the fossa were identified. The ganglion rootlets were severed, an incision was made to the dura, and the TG was removed from the petrous ridge using the internal carotid to reflect the dura and ganglion up and lateral to the skull base. In preparation for OCT mounting, blood clots were removed, and the internal carotid was trimmed adjacent to the V1 branch. The TG was directly mounted flat in OCT to maintain V1, V2, and V3 branch orientation. Tissue was then frozen and stored at -80 °C. Cryosectioning was performed as described above with 14-µm-thick slices placed onto coverslips coated with poly-D-lysine and fiducial beads. Slices were dried for 2 hours at -20 °C inside the cryostat, and then fixed in 4% PFA in 1× PBS for 20 minutes at 4 °C. The tissue was washed twice with 1× PBS and permeabilized overnight in 70% ethanol at 4 °C.

### Encoding probe hybridization

smFISH or MERFISH encoding probes were hybridized as described previously^12,43^. Briefly, cell culture samples were washed 3 times in 2× saline-sodium citrate (SSC; Thermo, AM9765) and once in 30% (v/v) formamide (Thermo, AM9342) in 2× SSC for 5 minutes at room temperature. Unless otherwise noted, samples were hybridized by incubation in a buffer containing smFISH or MERFISH probes at the reported concentrations suspended in 30% (v/v) deionized formamide, 10% (v/v) dextran sulfate (VWR, 97062-828), and 1 mg/mL yeast tRNA (Thermo, 15401029) in 2× SSC. In addition to encoding probes, this hybridization always included 1 µM of ‘anchor’ probes—30 nucleotides of poly dT mixed with locked-nucleic-acid dT and terminated with an acrydite moiety—to facilitate anchoring polyadenylated RNA to polyacrylamide gels as previously described^44^. A piece of parafilm (Sigma, P7793) was placed on the bottom of large petri dish, a 50-100 µL drop of the above hybridization solution was placed on this parafilm layer, and the sample was gently inverted onto this droplet. The concentration of formamide in this buffer was modified to the values listed in Figure 1 for the experiments involving encoding probes with different lengths of target regions. Unless otherwise specified, all samples were hybridized for 24 hours at 37 °C. For samples in which the hybridization duration was varied, this time was increased to either 4 days or 7 days in total.

To perform a melting and annealing step, a steel block (McMaster, 8983K211) was placed in an oven preheated to 50, 60, 70, or 80 °C. The block was allowed to equilibrate to that temperature for at least 1 hour. U-2 OS or tissue samples were prepared as above but a thin layer of mineral oil (Sigma, M5904) was placed between the bottom of the Petri dish and the steel block to promote thermal contact. As we found the parafilm partially melted at 70 °C, smFISH samples were instead placed upright in a petri dish and the droplet of hybridization solution as placed directly on them. After 1 hour at this elevated temperature, the samples and the steel block were moved to a 37 °C oven and incubated for 24 hours. As parafilm is generally intact at 60 °C, MERFISH samples at this temperature were placed cell-side down over 50 µL of hybridization solution and allowed to anneal and hybridize as described above.

After hybridization, samples were washed twice for 30 minutes each in 30% v/v formamide in 2× SSC at 47 °C. The exception was the samples prepared to explore the effect of target region length on encoding probe hybridization efficiency. These samples were instead washed at 47 °C in the same concentration of formamide in which they were hybridized.

Encoding probes associated with target region optimization experiments were stained with a total probe concentration of 2 µM with a 24-hour hybridization. These samples were stained for readout probes as described below using an RS0015-Cy5 probe that did not contain a disulfide bond. Hybridization duration measurements were conducted with the 130-RNA U-2 OS library encoding probes stained at the listed concentrations with hybridizations of the listed durations. Melting and annealing measurements were conducted with the 130-RNA U-2 OS library stained at total concentration of 4 µM. When MERFISH measurements were not conducted, these samples were stained with an RS0015-Cy5 and a RS0083-AF750 probe that did contain a disulfide bond. Adenine-spacer measurements were conducted with 1 µM of total probe with a hybridization duration of 24 hours. Readout reagent stability measurements and image buffer optimization measurements were conducted with a probe set targeting FLNA stained at total probe concentration of 1 µM for 24 hours. Both of these samples were stained with RS0015-Cy5 and RS0083-AF750 probes that did contain a disulfide bond.

Swiss roll MERFISH measurements performed with the old protocol followed the fixed-frozen MERFISH protocol described previously^43^. Briefly, following ethanol permeabilization samples were first stained for 24 hours at 37 °C with 2 µM anchor probes. Samples were then washed in 30% (v/v) formamide in 2× SSC, gel-embedded and digested (as described below). Samples were then hybridized with 35 µM of encoding probes over 3 days at 37 °C and finally washed and stored at 4 °C in 2× SSC until imaging. For the optimized protocols, Swiss rolls post-ethanol-permeabilization were stained with a hybridization mixture containing 2 µM of anchoring probes and 35 µM of encoding probes and immediately annealed at 60°C for 1 hour and then moved to a 37 °C oven and incubated for 24 hours similarly the protocol described for cell culture above. Samples were then gel embedded, digested, washed and stored at 4 °C in 2× SSC until imaging (as described below). Similarly, for the mouse samples used in readout screening, tissue slices were first stained with 1 µM anchor probes for 24 hours at 37 °C following ethanol permeabilization. The samples were then washed in 30% (v/v) formamide in 2× SSC, gel-embedded, digested, and stored in 2× SSC at 4 °C until imaging.

### Gel embedding and digestion

Cell culture or tissue samples were embedded in polyacrylamide gels and cleared as previously described^12,43^. Briefly, coverslips were washed briefly with a gel solution containing 4% 19:1 acrylamide/bisacrylamide (BioRad, 1610144), 0.15% (v/v) tetramethylethylenediamine (TEMED; Sigma, T7024), 0.03% (w/v) ammonium persulfate (Sigma, 09913), 300 mM NaCl (Thermo, AM9759), and 50 mM Tris-HCl (Fisher 15568-025) and then inverted onto a 50-100 µL droplet of the same solution placed on a GelSlick (Lonza, 50640) coated microscope slide. For cell culture, gel solutions also contained a 1:200,000 dilution of orange fiducial beads to serve as fiducial markers for MERFISH. The gel was allowed to polymerize for 2 hours.

Cell culture samples were then digested in a solution containing 0.8 M guanidine (Thermo, 24115), 0.5% (v/v) Triton-X, 5% (v/v) Tween-20 (Promega, H5152), 1% (v/v) proteinase K (NEB, P8107S), 30 mM EDTA (Thermo, AM9262), and 50 mM Tris-HCl for 12 hours at 37 °C. Tissue samples were digested in a buffer comprising 2% (v/v) sodium dodecyl sulfate (SDS; Thermo, AM9822), 1% (v/v) proteinase K, and 0.25% (v/v) Triton-X in 2×SSC for 1 day at 37 °C. This digestion was then repeated with fresh buffer for a second day. After digestion, samples were washed in 2× SSC for a total of five times at room temperature. Samples were used immediately or stored in 2× SSC at 4 °C until imaging.

### Imaging buffer compositions

To explore the role of buffer pH on the performance of imaging reagents, we used the following imaging buffers. All imaging buffers comprised 50 µM Trolox-quinone (made as described in Refs. ^43,48^), 0.5 mg/mL Trolox (Abcam, ab120747), 0.2% (v/v) PCD (OYCAmericas, 46852004), and 5 mM protocatechuic acid (PCA; Sigma, 37580). The original imaging buffer had these components in 2× SSC. All other imaging buffers replaced the 2× SSC with 300 mM NaCl and 50 mM of Tris-HCl, Glycine (Sigma, 50046), or TAPS (Sigma, T5316). For the rapid bleaching experiments shown in Fig. 4, the PCD concentration was reduced to 0.1% (v/v). In many cases, NaOH (Sigma, 72068) was added to bring the buffer to the final listed pH. For the MERFISH measurements with the optimized protocols, we supplemented all Tris-based buffer with 5 mM EDTA.

### MERFISH imaging system

All samples were imaged on one of several custom-built fluorescence microscopes as described previously^34,35^. Briefly, these microscopes were constructed from an inverted microscope body (Nikon) with a 5-color Celesta light engine, a motorized stage (Marzhauser or Zaber) for X/Y positioning, an objective nano-positioner (Mad City Labs) for Z positioning, a 60× 1.4 NA PlanApo oil objective (Nikon), and a camera system. This camera system consisted either of two sCMOS cameras (Orca Flash; Hamamatsu) with a camera splitter (Cairn) or a single CMOS camera (FLIR Blackfly). The fluorophores AF750 and Cy5 were excited at 750 nm and 635 nm, respectively. The orange fiducial beads were excited at 545 nm and DAPI was excited at 405 nm.

An automated flow system was used to coordinate delivery and incubation of imaging, cleavage, or hybridization buffers to the sample. This system consists of a flow chamber (Bioptechs, FCS2), a series of computer-controlled valves (Hamilton, MVP4), and a peristaltic (Gilson, F155004) or syringe (Hamilton, PSD4) pump. The microscope and fluidics system were controlled by open-source software (github.com/ZhuangLab/storm_control).

### smFISH and MERFISH imaging

smFISH and MERFISH measurements were performed as described previously^42,43^ with modifications only to buffer compositions. Briefly, for most samples, readout probes or the first set of readout probes were hybridized prior to loading the sample into the microscope and fluidics system. This ‘on-bench’ staining comprised hybridizing the samples in a readout hybridization buffer composed of 10% ethylene carbonate (Sigma, E26258) and 0.1% Triton-X in 2× SSC pH 7 supplemented with 3 nM of the desired readout probes for 15 minutes at room temperature. The sample was then washed in the same buffer but without readout probes and often with 3 µg/mL DAPI (Thermo, D1306) to stain nuclei for 10 minutes at room temperature. Samples were then immediately imaged. The optimized protocols replaced the 2× SSC with 300 mM NaCl, 50 mM TrisHCl, and 5 mM EDTA pH 8.

The imaging buffers are described above. Cleavage buffer was comprised of 50 mM TCEP (Goldbio, TCEP25) in 2× SSC (old protocols) or in 300 mM NaCl, 5 mM EDTA, and 50 mM Tris-HCl pH 8 (new protocols). Readout wash buffer was comprised of 10% ethylene carbonate and 0.1% Triton-X in 2× SSC (old protocols) or 300 mM NaCl, 5 mM EDTA, and 50 mM Tris-HCl pH 8 (new protocols).

During a MERFISH measurement, samples were stained by flowing 1.5 mL of each readout hybridization buffer containing 3 nM of the appropriate readout probes across the sample and then incubating the sample for 14 minutes. The sample was then washed with 0.75 mL of readout wash buffer and incubated in this buffer for 6 minutes. After imaging, the fluorophores attached to readout probes via a disulfide bond were removed by flowing 3 mL of cleavage buffer across the sample and then incubating for 15 minutes. Excess cleavage buffer was then removed by either washing the sample with 1.5 mL of 2× SSC for 3 minutes (old protocols) or with the same amount of 300 mM NaCl, 5 mM EDTA, and 50 mM Tris-HCl pH 8 for the same time (new protocols).

Readout probes (Table S2) were synthesized by Biosynthesis, Inc. Some readout probes were synthesized without disulfides linking the oligonucleotide to the fluorophore. These probes were synthesized by IDT.

### Nuclei counting

When it was necessary to determine the number of RNA molecules seen per cell, we estimated the number of cells by counting the number of nuclei in the sample. DAPI-stained samples were imaged with a 405-nm laser. To identify nuclei, the intensity of these images was used to create a segmentation mask with Otsu’s method defining the intensity threshold associated with pixels within nuclei. Standard morphological operations were then used to fill any holes within nuclei and to slightly erode these masks to disconnect the few nuclei that were close enough that their masks were touching. The watershed method was then applied to these binary masks to label individual masks as separate objects. The number of unique objects, i.e. nuclei, was used to estimate the number of cells per sample. The average RNA copy number per cell was determined by dividing the total number of RNA counts observed within each sample by the number of nuclei.

### Readout probe screening

Samples were prepared as described above for readout probe screening experiments. Namely, samples were prepared as fresh frozen blocks directly embedded in OCT after harvesting (mouse jejunum, ileum, colon, mouse and human trigeminal ganglia) or fixed directly after culture (U-2 OS, HEK, or HeLa cell lines). Samples were then cryosectioned (for tissue), dried at -20 °C for 2 hours, postfixed with 4% v/v PFA in 1× PBS at 4 °C for 20 minutes, washed twice in 4 °C 1× PBS, permeabilized with 70% ethanol, stained with 1 µM anchoring probes according to the encoding probe hybridization protocols, followed by gel embedding and clearing, and sample wash, all as described above. The only difference was that no encoding probes were included in the hybridization solution. Anchor probes were still included.

Samples were then run for MERFISH as described above with all listed readout probes in Fig. 4H included. Note that not all readout probes were screened across all sample types. Finally, the quality of readout probes was judged by visual inspection of mosaic images across large tissue or sample regions. More quantitative assessment is possible; however, we find it unnecessary as the problematic readout probes produce obvious fluorescence background.

### Optimized Buffer Compositions

For clarity, we provide here the composition of the final optimized buffers we recommend. The readout hybridization buffer contains 300 mM NaCl, 5 mM EDTA, 50 mM Tris-HCl, 10% ethylene carbonate, 0.1% Triton-X, and 3 nM of each readout probes. The readout wash buffer contains the same ingredients as above without the readout probes. The cleavage buffer contains 300 mM NaCl, 5 mM EDTA, 50 mM Tris-HCl, and 50 mM TCEP. The imaging buffer contains 300 mM NaCl, 5 mM EDTA, 50 mM Tris-HCl, 50 µM Trolox-quinone (made in water), 0.5 mg/mL Trolox, 0.2% (v/v) PCD, and 5 mM protocatechuic acid. The post-cleavage wash or rinsing buffer contains 300 mM NaCl, 5 mM EDTA, 50 mM Tris-HCl. All buffers should be pH 8 and may require adjustment with NaOH to reach this pH.

### Quantification of smFISH measurements

Properties of single-molecule signals, i.e., brightness and number, were determined via custom python scripts. Briefly, a high-pass gaussian filter was applied to remove low-intensity background. A binary mask was then created to remove extracellular fluorescence by eroding and dilating the existing smFISH spots within the cell. Remaining spots were localized and quantified through the application of a local maxima function. A noise-signal threshold was manually selected for each dataset to remove identification of single stray fluorophores or other low-intensity autofluorescence. This threshold was constant for all conditions within each channel for each separate dataset. Samples that examined the bleaching effect of fluorophores had probes localized and a noise-signal threshold identified from the first frame only, and successive frames recorded those localizations’ brightnesses and whether they met the noise-signal threshold.

### MERFISH image analysis

MERFISH data were decoded using a previously described pipeline^12,43^ (https://github.com/ZhuangLab/MERFISH_analysis) which was run on the Harvard Medical School Orchestra2 cluster. Background and foreground were selected in each measurement through a set of filters on the aggregate brightness of each RNA and of the number of pixels associated with RNA^12,43^.

### Readout reagent aging experiments

To determine the effect of aging imaging buffers, buffers were made as described above and either used immediately or stored in 50 mL falcon tubes under a ∼1”-thick layer of mineral oil to prevent the continuous consumption of oxygen by the scavenging systems. These buffers were stored at room temperature in the dark for 7 days, then a needle was used to extract the imaging buffer from underneath the mineral oil before use. To determine the aging of readout hybridization buffers, these buffers were made as described above and stored for the listed number of days at room temperature in the dark in the same plastic 24-deep-well cartridge (Fisher, NC1754429) used to hold these buffers in our automated fluid dispensing system. These cartridges were either placed in an open lab environment shielded from light, covered with parafilm, or covered via a thin layer of mineral oil placed on top of each buffer in the 24-deep-well cartridge.

### Calculation of the readout position and readout usage scores

To calculate the readout position score for individual barcodes, we assigned a numeric value to the hybridization round in which each bit was measured. We then summed the values corresponding to the locations in which a given barcode had a ‘1’. For example, the barcode ‘01 10 10 00 10’ would have a readout position score of 1+2+3+5=11 as it has a ‘1’ in the first, second, third, and fifth hybridization rounds.

To calculate the readout usage scores, we first calculated the abundance associated with each blank barcode that was measured. Then for each readout probe, we determine the readout usage score by adding the measured abundance of all blank barcodes that had a ‘1’ value in the bit that corresponds to that readout sequence and normalizing by the total counts observed for all blanks. To illustrate, let the abundance of blanks 1, 2, 3, and 4 be *N_1_*, *N_2_*, *N_3_*, and *N_4_*. If blank 1 and blank 3 used readout 1, then readout usage score for readout 1 would (*N_1_ + N_3_*)/(*N_1_*+*N_2_*+*N_3_*+*N_4_*).

## References

1. Zhuang, X. Spatially resolved single-cell genomics and transcriptomics by imaging. Nat. Methods 18, 18–22 (2021).

2. Moffitt, J. R., Lundberg, E. & Heyn, H. The emerging landscape of spatial profiling technologies. Nat. Rev. Genet. 23, 741–759 (2022).

3. Ke, R. et al. In situ sequencing for RNA analysis in preserved tissue and cells. Nature Methods 10, 857–860 (2013).

4. Gyllborg, D. et al. Hybridization-based in situ sequencing (HybISS) for spatially resolved transcriptomics in human and mouse brain tissue. Nucleic Acids Res. 48, e112 (2020).

5. Lee, H., Marco Salas, S., Gyllborg, D. & Nilsson, M. Direct RNA targeted in situ sequencing for transcriptomic profiling in tissue. Sci. Rep. 12, 7976 (2022).

6. Sountoulidis, A. et al. SCRINSHOT enables spatial mapping of cell states in tissue sections with single-cell resolution. PLoS Biol. 18, e3000675 (2020).

7. Wang, X. et al. Three-dimensional intact-tissue sequencing of single-cell transcriptional states. Science 361, aat5691 (2018).

8. Shi, H. et al. Spatial atlas of the mouse central nervous system at molecular resolution. Nature 622, 552–561 (2023).

9. Kalhor, K. et al. Mapping human tissues with highly multiplexed RNA in situ hybridization. Nat. Commun. 15, 2511 (2024).

10. Liu, S. et al. Barcoded oligonucleotides ligated on RNA amplified for multiplexed and parallel in situ analyses. Nucleic Acids Res. 49, e58 (2021).

11. Chen, K. H., Boettiger, A. N., Moffitt, J. R., Wang, S. & Zhuang, X. RNA imaging. Spatially resolved, highly multiplexed RNA profiling in single cells. Science 348, aaa6090 (2015).

12. Moffitt, J. R. et al. High-throughput single-cell gene-expression profiling with multiplexed error-robust fluorescence in situ hybridization. Proc. Natl. Acad. Sci. U. S. A. 113, 11046–11051 (2016).

13. Moffitt, J. R. et al. Molecular, spatial, and functional single-cell profiling of the hypothalamic preoptic region. Science 362, aau5324 (2018).

14. Xia, C., Fan, J., Emanuel, G., Hao, J. & Zhuang, X. Spatial transcriptome profiling by MERFISH reveals subcellular RNA compartmentalization and cell cycle-dependent gene expression. Proceedings of the National Academy of Sciences 116, 19490–19499 (2019).

15. Lubeck, E., Coskun, A. F., Zhiyentayev, T., Ahmad, M. & Cai, L. Single-cell in situ RNA profiling by sequential hybridization. Nature methods vol. 11 360–361 (2014).

16. Shah, S., Lubeck, E., Zhou, W. & Cai, L. In situ transcription profiling of single cells reveals spatial organization of cells in the mouse hippocampus. Neuron 92, 342–357 (2016).

17. Eng, C.-H. L. et al. Transcriptome-scale super-resolved imaging in tissues by RNA seqFISH. Nature 568, 235–239 (2019).

18. He, S. et al. High-plex imaging of RNA and proteins at subcellular resolution in fixed tissue by spatial molecular imaging. Nat. Biotechnol. 40, 1794–1806 (2022).

19. Janesick, A. et al. High resolution mapping of the tumor microenvironment using integrated single-cell, spatial and in situ analysis. Nat. Commun. 14, 8353 (2023).

20. Larsson, C., Grundberg, I., Söderberg, O. & Nilsson, M. In situ detection and genotyping of individual mRNA molecules. Nat. Methods 7, 395–397 (2010).

21. Femino, A M Fay, F S Fogarty, K Singer, R H. Visualization of Single RNA Transcripts in Situ. Science 280, 585–590 (1998).

22. Raj, A., van den Bogaard, P., Rifkin, S. A., van Oudenaarden, A. & Tyagi, S. Imaging individual mRNA molecules using multiple singly labeled probes. Nat. Methods 5, 877–879 (2008).

23. Shaffer, S. M., Wu, M.-T., Levesque, M. J. & Raj, A. Turbo FISH: A Method for Rapid Single Molecule RNA FISH. PLoS ONE 8, e75120 (2013).

24. Zhang, M. et al. Spatially resolved cell atlas of the mouse primary motor cortex by MERFISH. Nature 598, 137–143 (2021).

25. Bhattacherjee, A. et al. Spatial transcriptomics reveals the distinct organization of mouse prefrontal cortex and neuronal subtypes regulating chronic pain. Nat. Neurosci. 26, 1880–1893 (2023).

26. Allen, W. E., Blosser, T. R., Sullivan, Z. A., Dulac, C. & Zhuang, X. Molecular and spatial signatures of mouse brain aging at single-cell resolution. Cell 186, 194–208.e18 (2023).

27. Osterhout, J. A. et al. A preoptic neuronal population controls fever and appetite during sickness. Nature 606, 937–944 (2022).

28. Chen, R. et al. Decoding molecular and cellular heterogeneity of mouse nucleus accumbens. Nat. Neurosci. 24, 1757–1771 (2021).

29. Sun, W. et al. Spatial transcriptomics reveal neuron-astrocyte synergy in long-term memory. Nature 627, 374–381 (2024).

30. Fang, R. et al. Conservation and divergence of cortical cell organization in human and mouse revealed by MERFISH. Science 377, 56–62 (2022).

31. Zhang, M. et al. Molecularly defined and spatially resolved cell atlas of the whole mouse brain. Nature 624, 343–354 (2023).

32. Yao, Z. et al. A high-resolution transcriptomic and spatial atlas of cell types in the whole mouse brain. Nature 624, 317–332 (2023).

33. Lu, Y. et al. Spatial transcriptome profiling by MERFISH reveals fetal liver hematopoietic stem cell niche architecture. Cell Discov. 7, 47 (2021).

34. Watson, B. R. et al. Spatial transcriptomics of healthy and fibrotic human liver at single-cell resolution. Nat. Commun. 16, 319 (2025).

35. Petukhov, V. et al. Cell segmentation in imaging-based spatial transcriptomics. Nat. Biotechnol. (2021) doi:10.1038/s41587-021-01044-w.

36. Cadinu, P. et al. Charting the cellular biogeography in colitis reveals fibroblast trajectories and coordinated spatial remodeling. Cell 187, 2010–2028.e30 (2024).

37. Farah, E. N. et al. Spatially organized cellular communities form the developing human heart. Nature 627, 854–864 (2024).

38. Choi, J. et al. Spatial organization of the mouse retina at single cell resolution by MERFISH. Nat. Commun. 14, 4929 (2023).

39. Sarfatis, A., Wang, Y., Twumasi-Ankrah, N. & Moffitt, J. R. Highly multiplexed spatial transcriptomics in bacteria. Science 387, eadr0932 (2025).

40. Nobori, T. et al. Time-resolved single-cell and spatial gene regulatory atlas of plants under pathogen attack. bioRxiv (2023) doi:10.1101/2023.04.10.536170.

41. Long, K. A., et al. Spatial transcriptomics reveals expression gradients in developing wheat inflorescences at cellular resolution. bioRxiv (2024) doi:10.1101/2024.12.19.629411.

42. Moffitt, J. R. & Zhuang, X. RNA imaging with multiplexed error-robust fluorescence in situ hybridization (MERFISH). Methods Enzymol. 572, 1–49 (2016).

43. Cadinu, P. et al. Imaging the intestinal transcriptome with multiplexed error-robust fluorescence in situ hybridization (MERFISH). Curr. Protoc. 5, e70111 (2025).

44. Moffitt, J. R. et al. High-performance multiplexed fluorescence in situ hybridization in culture and tissue with matrix imprinting and clearing. Proc. Natl. Acad. Sci. U. S. A. 113, 14456–14461 (2016).

45. Dey, S. et al. DNA origami. Nat. Rev. Methods Primers 1, (2021).

46. Rasnik, I., McKinney, S. A. & Ha, T. Nonblinking and long-lasting single-molecule fluorescence imaging. Nat. Methods 3, 891–893 (2006).

47. Aitken, C. E., Marshall, R. A. & Puglisi, J. An oxygen scavenging system for improvement of dye stability in single-molecule fluorescence experiments. Biophys. J. 94, 1826–1835 (2007).

48. Cordes, T., Vogelsang, J. & Tinnefeld, P. On the mechanism of Trolox as antiblinking and antibleaching reagent. J. Am. Chem. Soc. 131, 5018–5019 (2009).

49. Bull, C. & Ballou, D. P. Purification and properties of protocatechuate 3,4-dioxygenase from Pseudomonas putida. A new iron to subunit stoichiometry. J. Biol. Chem. 256, 12673–12680 (1981).

50. Patil, P. V. & Ballou, D. P. The use of protocatechuate dioxygenase for maintaining anaerobic conditions in biochemical experiments. Anal. Biochem. 286, 187–192 (2000).

51. Swoboda, M. et al. Enzymatic oxygen scavenging for photostability without pH drop in single-molecule experiments. ACS Nano 6, 6364–6369 (2012).

52. Walz, S. et al. Activation and repression by oncogenic MYC shape tumour-specific gene expression profiles. Nature 511, 483–487 (2014).

53. Lin, M. T.-Y. et al. Culture of primary neurons from dissociated and cryopreserved mouse trigeminal ganglion. Tissue Eng. Part C Methods 29, 381–393 (2023).

