## Supplementary Figures and Captions for "Protocol Optimization Improves the Performance of Multiplexed RNA Imaging"

### Supplementary Information

#### Supplementary Figures

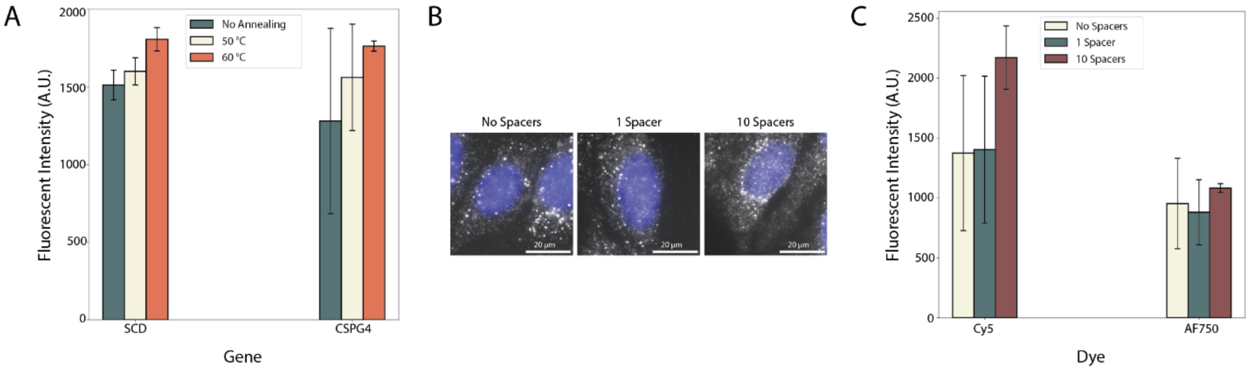


**Supplementary Figure 1. Properties of the Design and Hybridization of Encoding Probes that May Improve Signal Brightness.** (**A**) Single-molecule signal brightness for encoding probes targeting SCD or CSPG4 stained using a protocol without melting and annealing (No Annealing) or with an initial melting step and an annealing step that begins at the listed temperature (Methods). (**B**) Images of U-2 OS cells stained with encoding probes that target SCD and which have two readout sequences separated by 0, 1, or 10 adenines (spacers). Gray: mRNA signal. Blue: DAPI. Scale bars: 20 µm. (**C**) Single-molecule signal brightness for encoding probes targeting SCD with different numbers of adenine spacers as in (B) for the readout probes conjugated to Cy5 or AF750. For A and C: Bars and error bars represent the average or standard deviation across three biological replicates of the average molecular brightness seen within each replicate.

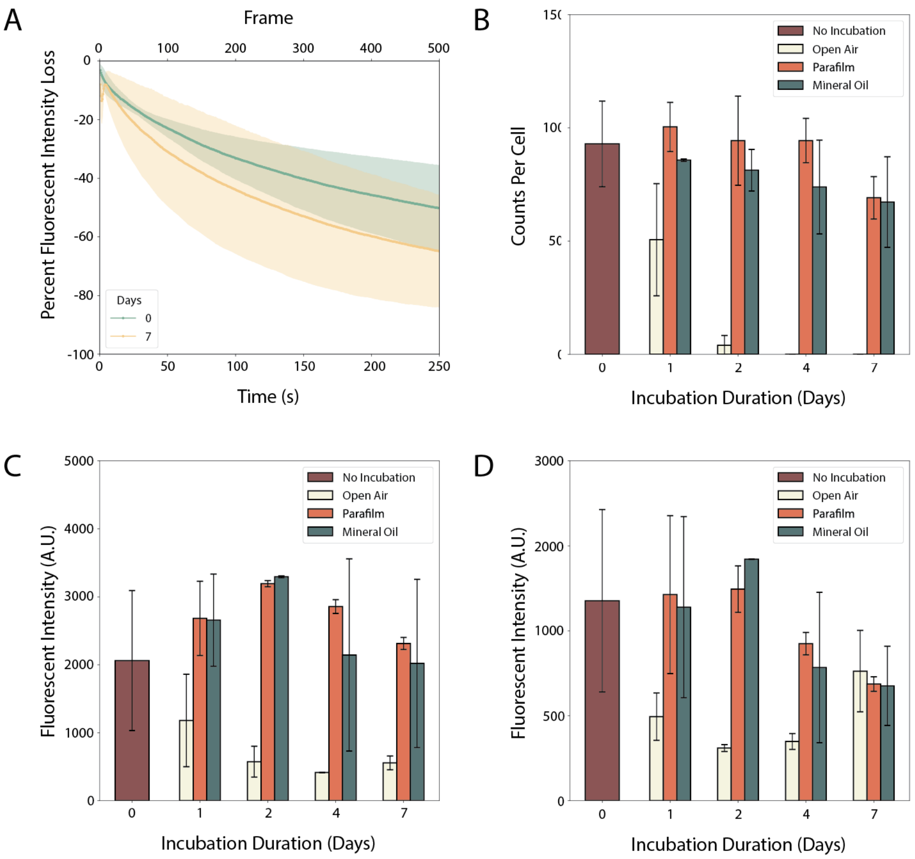


**Supplementary Figure 2. Additional Measurements of the Aging of MERFISH Readout Reagents.** (**A**) Single-molecule signal brightness measured for the AF750 channel versus the total exposure time for samples as in (Fig. 2C). Intensity is measured in precent brightness decrease relative to that observed in the first image. (**B**-**D**) The number of RNA molecules observed per cell in the AF750 channel (B) or the average molecular brightness for the Cy5 (C) or AF750 (D) channels. No incubation indicates reagents prepared freshly, ‘Open Air’ indicates reagents aged in the dark but exposed to the room environment, ‘Parafilm’ indicates reagents aged in the dark but protected from the room environment with parafilm, and ‘Mineral Oil’ indicates reagents aged in the dark and protected from the room environment and oxygen by a layer of mineral oil. For A, B, C, and D: Solid lines or bars represent averages while shaded areas or error bars represent the standard deviation of the plotted values observed across two replicates.

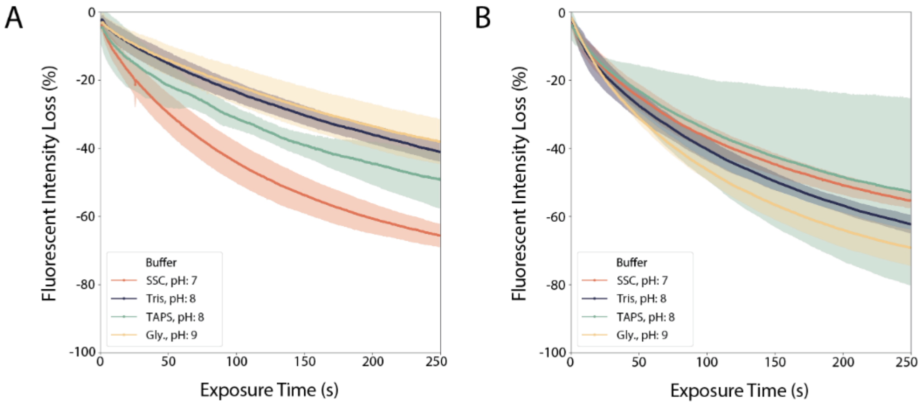


**Supplementary Figure 3. The Photostability of Cy5 and AF750 in Different Imaging Buffers.** (**A**,**B**) The average signal brightness measured for individual molecules in the Cy5 (A) or AF750 (B) channel versus the total exposure time. Intensity is measured in the precent decrease in brightness relative to that observed in the first image. The solid line represents the average across all molecules in two biological replicates while the shaded areas represent the standard deviation observed around this average. These measurements were conducted several minutes after imaging buffer was introduced into the sample to allow the scavenging system to reduce the oxygen levels to steady state.

### Supplementary Table Captions

**Supplementary Table 1. Oligonucleotide Sequences Used in smFISH Experiments. (Provided as a separate xlsx file).** The 'Target' column contains the name of the targeted mRNA. The 'Experiment' column contains a short description of the experiment for which the encoding probe was designed. The 'Sequence' column contains the sequence of the encoding probe.

**Supplementary Table 2. MERFISH Readout Probe Sequences. (Provided as a separate xlsx file).** The 'Name' column contains a unique name for each readout probe. The 'Sequence' column contains the sequence of the readout probe. 'Cy5-S-S-' and 'AF750-S-S-' indicate that a Cy5 or an AF750 was conjugated to the probe via a disulfide bond, respectively.

**Supplementary Table 3. MERFISH Encoding Probe Sequences for a Mouse Gastrointestinal Tract Library (Provided as a separate xlsx file).** The 'Name' column contains a unique name for each encoding probe template molecule. The 'Sequence' column contains the sequence of the encoding probe template.

**Supplementary Table 4: MERFISH Codebook for a Mouse Gastrointestinal Tract Library (Provided as a separate xlsx file).** In the first sheet, the 'Gene' column indicates the targeted gene, the 'ID' column indicates a unique ID for the targeted isoform, and the 'Barcode' column lists the binary barcode assigned to each gene. An entry of 'Blank' indicates a blank barcode. In the second sheet, the 'Bit' column lists the bit position, and the 'Name' column lists the name of the readout sequence (as in Table S2) used to readout that bit value.
